# Cross-reactive anti-hemagglutinin (HA) IgG responses are shaped by previous long-interval monovalent H5 vaccination and highly correlated with HA antigenic distance

**DOI:** 10.1101/2020.10.14.340448

**Authors:** Jiong Wang, Dongmei Li, Sheldon Perry, Shannon P. Hilchey, Alexander Wiltse, John J. Treanor, Mark Y. Sangster, Martin S. Zand

**Affiliations:** Department of Medicine, Division of Nephrology, University of Rochester Medical Center, Rochester, NY USA; Informatics Core, Clinical and Translational Science Institute, University of Rochester, Rochester, NY US; Department of Medicine, Division of Infectious Diseases, University of Rochester Medical Center, Rochester, NY USA; Department of Immunology, Vaccine Center, University of Rochester Medical Center, Rochester, NY USA; Rochester Center for Health Informatics, University of Rochester Medical Center, Rochester, NY USA

**Keywords:** H5 monovalent influenza vaccine (MIV), antigenic distance, hemagglutinin(HA), antibody landscapes, Original antigenic sin (OAS), HA imprinting

## Abstract

Avian H5 influenza is an emerging influenza strain with the potential for human pandemic spread. One unresolved issue in pandemic vaccine preparedness is to what extent a vaccine recall response depends on the interval between the priming and boosting vaccinations. In this study, we analyzed the anti-H5 HA IgG responses to an H5 A/Indonesia/5/2005 boosting vaccination in three cohorts: (1) a short interval boosting cohort that received a prime and boost 28 days apart, (2) a long-interval boosting cohort that received an H5 A/Vietnam/203/2004 priming vaccination 5 years before boosting, and (3) a double long-boost cohort that received single doses of all three vaccines separated by 5-6 year intervals. Anti-HA IgG levels were measured using a multiple-plex assay against 21 H5 and 16 seasonal strains covering both influenza phylogenetic groups. We used the antigentic distance between the vaccine strain and each HA in the assay panel to define the antibody response landscape. Both single and double long-interval boosting with the H5 variant vaccine elicited a broad antibody response to all H5 subtype strains, and double boosting resulted in sustained, vaccine-specific, anti-HA IgG levels over a six month period. Antibody-mediated immune responses were shaped by prior H5 exposure history, and the magnitude of both vaccine specific and cross-reactive anti-H5 HA IgG responses was highly correlated with the relative antigenic distance between the measured and the vaccine HAs. We conclude that the relative antibody landscape method can be used to quantify the phenomenon of antigenic imprinting on human influenza vaccine immune responses.

**IMPORTANCE:** A signi1cant obstacle to development of a universal influenza vaccine is understanding the relationship between multidimensional host humoral immunity, prior antigen exposure, and viral antigenicity. In this study, we used a multiplexed antibody assay to measure antibody cross-reactivities against antigenically similar H5 influenza virus strains. This work uses a novel method, relative antibody landscapes, to analyze the relationship between immune response and antigenic distance between the target H5 vaccine HA and other HAs in the assay after a boosting vaccination. This method improves analysis of immune responses by relating antigen exposure history to the influenza vaccination antibody response. This study also revealed that multiple vaccine boosting over several years can generate high levels of long-lasting cross-reactive antibodies against the priming H5 strains vaccine that subjects received, suggesting the HA imprinting mechanism(s) have a strong influence in the adult antibody response to H5 MIV vaccination.

## INTRODUCTION

Seasonal influenza (flu) virus infection is a significant global health threat, infecting over millions annually with significant deaths. The primary protection against influenza infection comes from antibodies directed against the influenza hemagglutinin (HA) protein, a major vaccine target(1). Pandemic influenza infections occur when new influenza viruses emerge with significant mutations in HA and evade prior influenza immunity. HA is composed of two domains, the highly plastic globular HA1 head domain and the conserved HA2 stalk domain. The hypervariable head domain of HA is believed to be immunodominant. Infection and vaccination primarily elicit strain-specific neutralizing antibodies, with limited cross-reactivity to divergent strains. In contrast, antibodies that target epitopes on the stalk domain can broadly cross-react with multiple influenza strains (2).

A number of influenza strains have caused pandemics after animal to human transmission, particularly the H5 strain known to circulate in poultry(3, 4). The first human H5N1 infection was reported in 1997 during a poultry H5 outbreak in Hong Kong (5). From 2003 to January 2015, a total of 694 laboratory-confirmed human H5 cases were reported from 16 countries with 58% mortality rate (6). Almost all human cases have occurred as a direct result of close contact with infected birds or chickens. In 2015 a sizeable H5N2 outbreak within turkey and chicken farms occurred in the US, but were not transmitted to humans (7). However, the H5 virus has significant potential to cause future human influenza pandemics given its high mutation and recombination rates(8). Currently, there are no commercial human H5 vaccines available. in addition, H5 non-adjuvanted monovalent influenza vaccine (MIV) generally requires a prime and boost strategy (9, 10), as the primary anti-HA antibody response is very low (11, 12, 13, 14, 15). Thus, understanding the effects of prior immunologic memory, cross-reactive immunity, and the emergence of broadly cross reactive IgG mediated immunity are critical to effective vaccine development.

One fundamental hurdle to eliciting effective immune responses against emerging influenza strains is the concept of "original antigenic sin" (OAS), variously referred to now as HA imprinting (16), or HA seniority (17, 18, 19). When a person is sequentially exposed to two related influenza virus strains, they tend to elicit an immune response dominated by antibodies against the first strain they were exposed to(20, 21). This is true even following a secondary infection or vaccination. Thus, the immune response to a new influenza viral infection or vaccination is at least partially shaped by preexisting influenza immunity. For development of vaccines against antigenically dissimilar influenza strains, it is critical to understand the antibody response against antigenically similar virus stains and vaccine development, especially within the context to OAS.

Based on the phylogenetic distance of HA genes, ten clades of H5 HA (clade 0-9) have been identified within the H5N1 virus subtype (22). H5N1 viruses from clades 0, 1, 2, and 7 have the capacity to infect humans (23), and a universal H5 influenza vaccine would be able to induce broad cross-reactivity, against all of these clades. However, there are three distinct antigenic clusters, as determined by antigenic cartography generated with neutralizing antibody levels induced by H5 HA DNA vaccination in mice (3). This suggests the possibility that HA imprinting may impede generation of broadly cross-reactive H5N1 antibodies if the prime and boost H5N1 vaccine strains reside in different antigenic clusters. To address this issue, this study examines the effect of antigenic distance between a primary H5 vaccine strain and a subsequent, long term vaccine boost strain in human subjects.

In this manuscript, we re-evaluated serum samples from a previous H5 human vaccine study (DMID 08-0059)(24) using multi-dimensional measurement of anti-H5N1 HA IgG reactivities. Samples were collected during the inactivated A/Indonesia/5/05 (Ind05) MIV study from subjects had received two primed H5 MIV (A/Hong Kong/156/97 (HK97) in 1997–1998 and A/Vietnam/1203/04 (VN04) in 2005–2006), with one VN04 primed and naive control groups. Anti-HA antibody responses were measured by mPlex-Flu assay(25) to simultaneously evaluate the magnitude and breadth of the IgG repertoire directed against HAs from 21 H5 influenza virus strains and 9 other IAV strains (H1, H3 H7, H9). We hypothesized that as the relative antigenic distance between the original priming and the new H5 boosting vaccine strain becomes smaller (i.e. the strains are more antigenically similar), the greater the increase in the anti-HA IgG response to original H5 MIV strain. Thus, in a vaccine response, the original HA imprinting influences much later vaccine responses. We discuss the relevance of these findings to the development of influenza vaccines that induce broad antibody-mediated protection (i.e. universal influenza vaccines).

## RESULTS

### Characteristics of subjects

Prior exposure to the predominant seasonal H1 or H3 influenza strain circulating close to a subject’s birth year can alter H5 or H7 infection and death rates (26, 27, 16). Thus, we first tested tested for differences in age that could alter the antibody levels between the H5 vaccine groups. To assess the birth year related influenza exposure history, we regrouped the study cohorts based on two key birth years: 1968 and 1977, when H3 and H1, respectively, became the dominant circulating influenza A strains (Table 1) (16). Subjects without baseline (pre-vaccination) serum samples were excluded, leaving a total of 55 subjects. The H5 naive subjects (Naive, *n* = 12) and primed subjects (L-boost, *n* = 30) previously received an inactivated subvirion influenza A/Vietnam/1203/04 (VN04) vaccine in 2005–2006(9). The double primed group (DL-boost, *n* = 13) received the recombinant influenza A/Hong Kong/156/97 vaccine (A/HK97) in 1997 - 1998 (11) and the VN04 vaccine in 2005 - 2006. We found no significant difference in birth year distributions between the cohorts (P > 0.05; Fisher’s exact test), suggesting that the effects of flu exposure history on the H5 MIV vaccine response should be similar across the three groups.

**TABLE 1.** The number of subjects stratified by birth year in each cohort of the DMID 08-0059 study. Subjects were grouped by birth year based on key years when either H3 or H1 representing the predominant circulating seasonal flu strains, as prior exposure history might influence the antibody responses to the H5 vaccines.

| Group | Birth Year |  |  | Total |
| --- | --- | --- | --- | --- |
|  | < 1968 | 1968 to 1977 | > 1977 |  |
| Naïve | 10 (83) | 1 (8) | 1 (8) | 12 |
| Long-Interval Boost (L-boost) | 24 (80) | 3 (10) | 3 (10) | 30 |
| Double long-Interval Boost (DL-boost) | 11 (85) | 2 (15) | 0 (0) | 13 |

### High anti-H5 IgG responses resulting from long-interval boosting are shaped by the priming vaccine strain

Using a 48-HA mPLEX-Flu assay panel, we observed that IgG response levels against the HA of A/Indonesia/5/05 (Ind05), VN04 and HK97 were very low in the naive group, and about two-fold higher in the short interval boosting (S-boost) group who were boosted after 28 days (FIG 2 A, B). In both primed groups (L-boost and DL-boost), however, inactivated Ind05 MIV induced ∼5-fold higher vaccine-specific antibody levels by 14 days post-vaccination. Anti-VN04 and HK97 IgG levels increased ∼7-8 fold, also peaking at 14 days in both primed groups (FIG 2). While both primed groups had higher pre-existing (day 0) anti-H5 IgG levels, their IgG response kinetic curves against the vaccine strains were similar. These differences result in a relative increase in the DL-boost group’s anti-HA antibody levels peaking at 3.5-fold (FIG2, D), even though the post-boosting IgG levels are similar in the S- and DL-boost groups. In both groups, anti-H5 HA antibodies levels remained high for over six months. These results are consistent with the previous 1nding that non-adjuvanted MIVs are poorly immunogenic in naive subjects (11, 12, 13, 14, 15), and long-interval boosting with H5 antigenic variant MIVs elicits significant and robust antibody responses (9, 24). However, this is the first report to show differences in antibody response induced by single vs. double long-interval MIV boosting.

**FIG 1.**
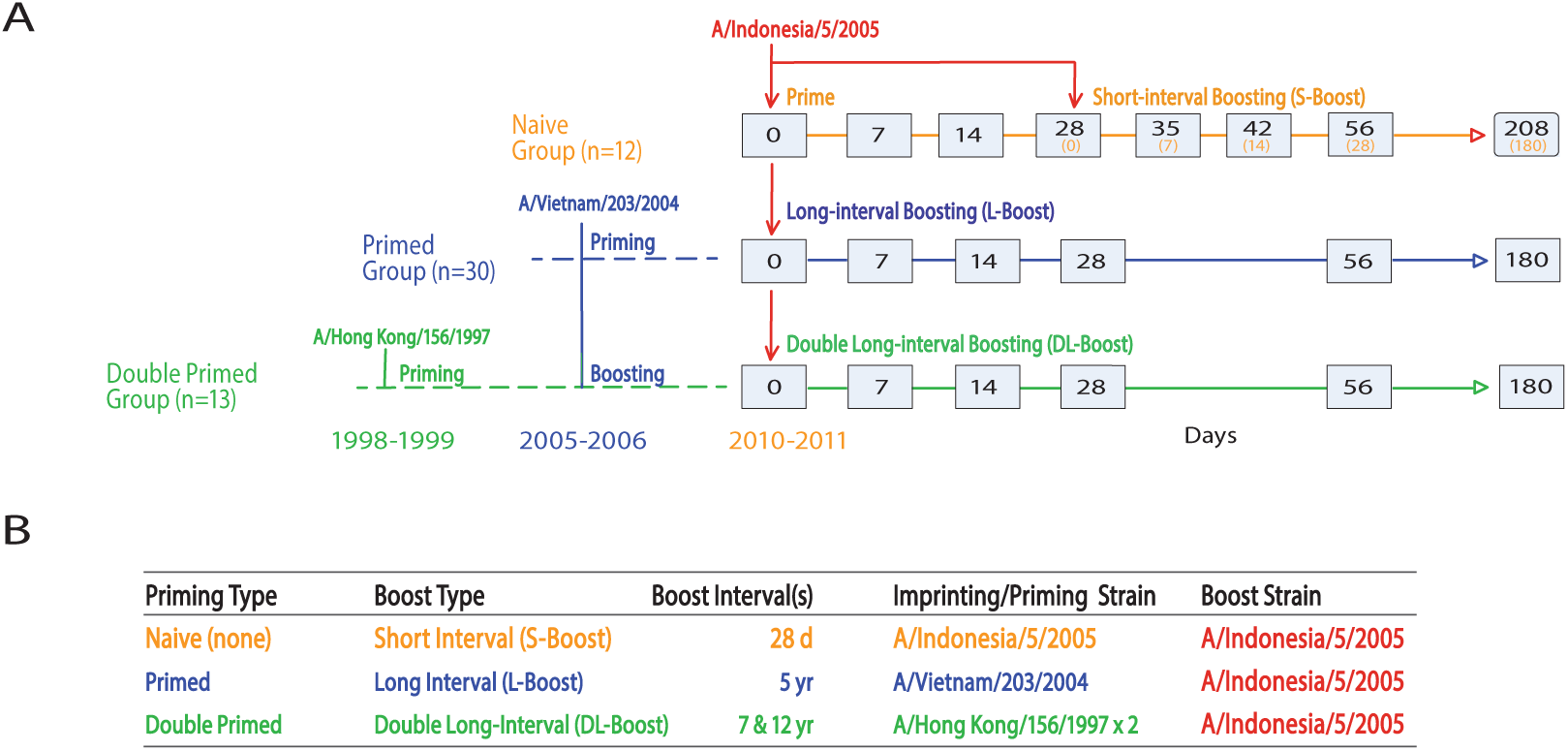
Vaccination strategy. (A) Trial and sampling design: All subjects in the DMID 08-0059 study cohorts were vaccinated with inactivated A/Indonesia/5/05 (Ind05) intramuscular influenza vaccine. The Naive group; S-boost received the Ind05 vaccine on day 0, and short interval boosting on day 28. The primed long-interval boost (L-boost) group had previously received the inactivated subvirion influenza A/Vietnam/1203/04 (Vie04) vaccine in 2005–2006; and the primed double long interval boost (DL-boost) group additionally received the baculovirus expressed recombinant influenza A/Hong Kong/156/97 vaccine (HK97) in 1997–1998. Both L-boost and DL-boost groups also received long-interval vaccination with Ind05 on the day 0. Grey boxes indicate serum sampling. A) Summary of prime and boost strains and groups.

**FIG 2.**
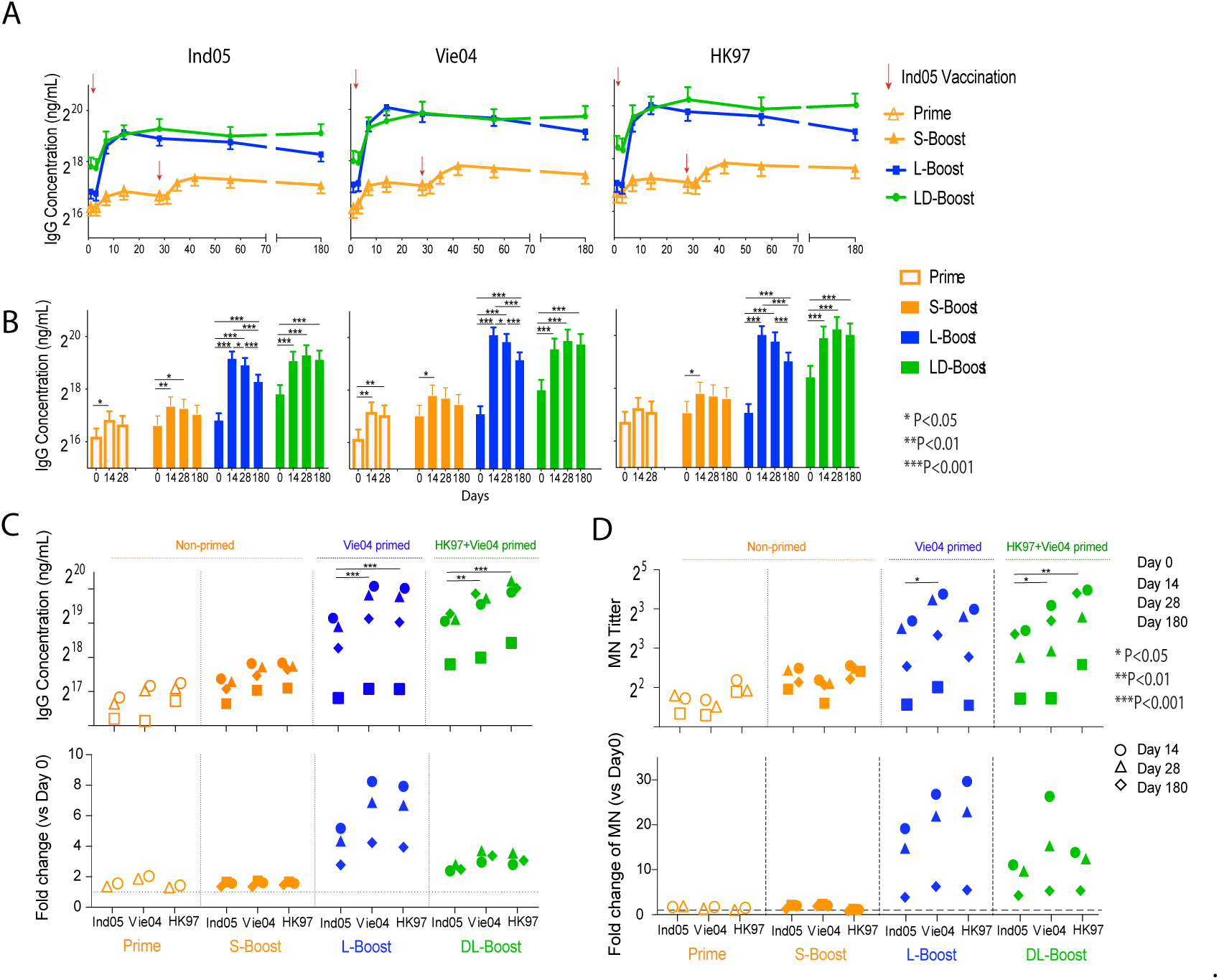
The effects of prior vaccination with H5 monovalent influenza vaccine (MIV) on multiplex HA antibody responses against three different H5 virus boosting vaccine strains. The mean and standard deviation of IgG concentration for each group were estimated by the mPlex-Flu assay. Antibody concentrations were adjusted within the linear mixed effects models using age at enrollment, gender, ethnicity (Caucasian vs. non-Caucasian), dose (two dose levels: 15 and 90 *µg*), and assay batch (five batches) (28, 29). A. The H5 kinetic antibody levels against three vaccine strains after MIV H5 vaccination with A/Indonesia/05/2005 (Ind05; clade 2). The primed response (Prime, unfilled symbols) and short-interval boost response (S-boost, filled symbols) of naive subjects; the long-interval boost response (L-boost) after one dose of Ind05 MIV in subjects primed by Vie04 MIV the 5 years previously, and in the subjects who were double primed with Vie04 (5 years previously) and A/Hong Kong/156/1997 (HK97; clade 0) HK97 (12-13 years previously), as the double long-interval boost response (DL-boost), against Ind05, A/Vietnam/1203/2004 (Vie04; clade 1) and A/Hong Kong/156/1997 (HK97; clade 0), three vaccine H5 strains. B. Comparison of antibody responses between time points in the same groups for each vaccine strain. C. The antibody concentrations against each vaccination strain and fold changes as compared to day 0, grouped by study cohort. D. The antibody titers for micro-neutralization (MN) against each vaccination strain and fold changes, as compared to day 0, grouped by study cohort. The original MN assay data was been re-analysed with linear mixed effects modeling, as above. Shown are the geometric mean of titers. * P<0.05, **P<0.01, ***P<0.001 Linear contrasts within the linear mixed effects model framework were used to conduct the statistical comparisons.

Importantly, we also found that the Ind05 MIV elicited robust antibody responses against the two previous priming H5 strains (VN04, HK97) in both vaccine groups, and that the anti-HA IgG responses shared similar kinetic patterns. Interestingly, Ind05 MIV elicited higher levels of IgG antibodies to VN04 and HK97 than to Ind05. In order to directly compare the effects of the priming virus strain, we plotted the concentrations of anti-H5 HA by groups, shown in FIG2 C, and the fold change of antibody concentrations against three vaccine strains of the different groups (FIG2 D). The results revealed higher antibody levels against the HA of VN04 in the L-boost group, and HK97 in the DL-boost group, which were the first H5 viral strains subjects were respectively vaccinated against. These results could be interpreted as indicative of HA imprinting (21, 20), in which subjects generate a robust antibody response against the H5 influenza strain they were first exposed to, by infection or vaccination, and maintain this response over their entire lifetime (30).

To confirm the protective activities of the higher level of long-lasting antibodies in the L-boost and DL-boost groups we re-analyzed the HAI and MN data in the DMID 08-0059 study using the generalized linear mixed effects models with identity link functions as we have previously described (28, 29). The results confirmed that all three H5 MIV strain vaccines induced serum with viral neutralizing capacity and could protect cells from viral infection (FIG 2 D and FIG S8).

### Relative antigenic response landscapes of H5 MIV HAs

Our results also raised another fundamental question: Does the magnitude of the imprinted recall response to the primoriginal H5 HA correlate with the antigenic distance between the HAs of the prime and boost strains? We hypothesized that the antigenic distance between the vaccine strain and a target H5 HA is inversely correlated with the cross-relativity of antibody response induced by the H5 MIV. In other words, smaller antigenic distances from the 1rst influenza strain (imprinting strain) produce larger IgG responses. To answer this question, we performed antigenic cartography to quantitatively evaluate the antigenic distances between H5 clades and subclades.

Recombinant H5 HA proteins were expressed and purified. Strains were chosen to cover all 10 H5 clades (0-9) or subclades, and 4 new H5 avian strains (Cl4.4.4.3) cloned in the US (TABLE S1, and FIG S1). Antibody reactivity to these strains was plotted against mouse anti-H5 HA IgG serum reactivity generated utilizing a monovalent DNA vaccination approach (FIG S2 A). We thus generated a comprehensive antigenic distance matrix between 17 H5 influenza strains and each of 21 H5 and 9 other influenza virus strains using the mPlex-Flu assay. The individual antibody levels against H5 viruses are shown as MFI units at specific dilutions, with the dilution factors being normalized using a generalized linear model with an identity link function for the sera samples. We used classical multidimensional scaling (MDS) method(31) to project relative distances between strains into 2 dimensions, and the matrix data was created by calculating the Euclidian distance matrix from two-dimension coordinates. Finally, we used a modification of the approach of Smith, et al.(32) to visualize the antigenic distance between influenza virus HAs(32, 3) (FIG S2, C). This approach accounts for the continuous nature of the mPlex-Flu assay data and the consistent range of estimated strain-specific binding(28, 29), yielding the same results as antigenic cartography. The antigenic distance matrix was also generated from the above multiplex data of mPlex-Flu assay using the DNA vaccine anti-sera (FIG S3).

In order to show the relative antigenic distance between individual HAs and the H5 MIV strains (FIG 3 B), we plotted the distance of each H5 HA relative to the 3 vaccine strains: HK97 (X-axis), Vie04 (Y-axis) and Ind05 (Z-axis). Each marker diameter represents the magnitude of the IgG concentration 14 days after MIV boosting. This allowed visualization of the magnitude of the antibody response against specific H5 HAs, associated with antigenic distance in different cohort groups with respect to both prime and boost vaccine strains. The same diagram allowed visualization of H5 strain vaccine strain relative distances from other H5 strains. Naive subjects had low anti-HA IgG levels against all H5 strains after priming and short-interval boosting with MIV. However, the L-boost and DL-boost groups had significantly enhanced antibody responses after 14 days, with higher IgG responses to H5 strains in the Vie04 and HK97 cluster groups than to the viruses in the MIV Ind05 cluster group, which are antigenically similar to the strain of the more recent MIV (FIG 3 C). These data more clearly show the relationship between the anti-HA IgG antibody response and the antigenic distances to the reference strains: higher cross-reactive antibody levels are elicited against the HAs from strains in the same cluster group with the first priming virus strain.

**FIG 3.**
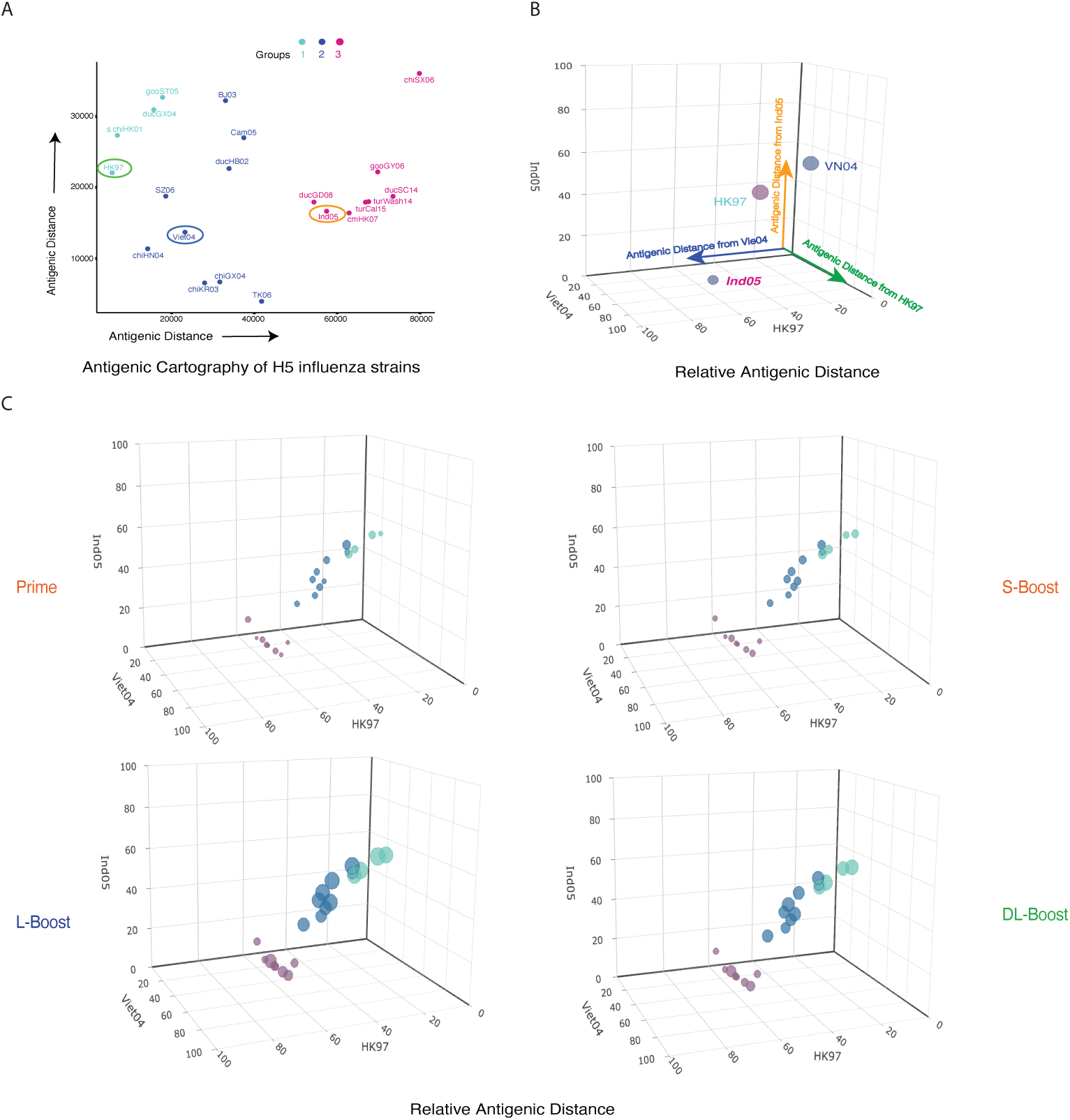
HA antibody responses were plotted against the related antigenic distances to each monovalent H5 vaccination (MIV) strain of different study cohorts. A. Antigenic cartography of 21 H5 influenza virus strains generated by mPlex-Flu assay of antisera against 17 anti-H5 influenza viruses, and plotted using using classical multi-dimensional scaling (MDS;see Methods). The three vaccine strains are circled. B. We then used three dimensional plots to show the relative antigenic distance of all mPLEX-Flu target HAs to the three vaccine strains A/Hong Kong/97 (HK97, clade 0), A/Vietnam/1203/2004 (VN04; clade 1), A/Indonesia/05/2005 (Ind05; clade 2). C. The IgG response of subjects in the DMID 08-0059 study to 21 H5 strains plotted in 3D bubble plots. The relative antigenic distances of the 21 H5 strains assayed were plotted against their antigentic distance to each of the three MIV strains to determine giving 3D-antigenic cartography. The bubble size represents the concentration (10^4^ng/mL) of IgG against an H5 influenza strain at day 14 post MIV boosting. (A) Using unsupervised hierarchical clustering, three H5 antigenic groups were identified. Interactive 3D bubble plots can be accessed through the following links: Prime group (http://rpubs.com/DongmeiLi/565996); S-boost group(http://rpubs.com/DongmeiLi/565998); L-boost: (http://rpubs.com/DongmeiLi/565989); DL-boost: (http://rpubs.com/DongmeiLi/565994).

### Long-interval boosting of MIV elicited heterogeneous IgG responses against all H5 clade/subclades, which were correlated with the antigenic distance to the 1rst primed virus strains

We next generated antigenic landscape plots (27) to visualize the magnitude of serological responses in relation to the antigenic distance between the vaccine strain HA and the H5 HAs in the mPlex-Flu panel. We 1rst focused on the relationship between the magnitude of boosted IgG response and the antigenic distance between the boost HA and the three H5 vaccine strains. To this end, IgG antibody concentrations against 21 H5 strains were measured by mPLEx-Flu assay for each cohort on days 9, 14, and 28, which were plotted against their relative antigentic distances to Ind05 (FIG 4A, B), Viet04 (FIG 4C, D), and HK97 (FIG 4E, F). Correlation test results are given in the 1gure inset, and all data are presented in FIG S4, S5, S6).

**FIG 4.**
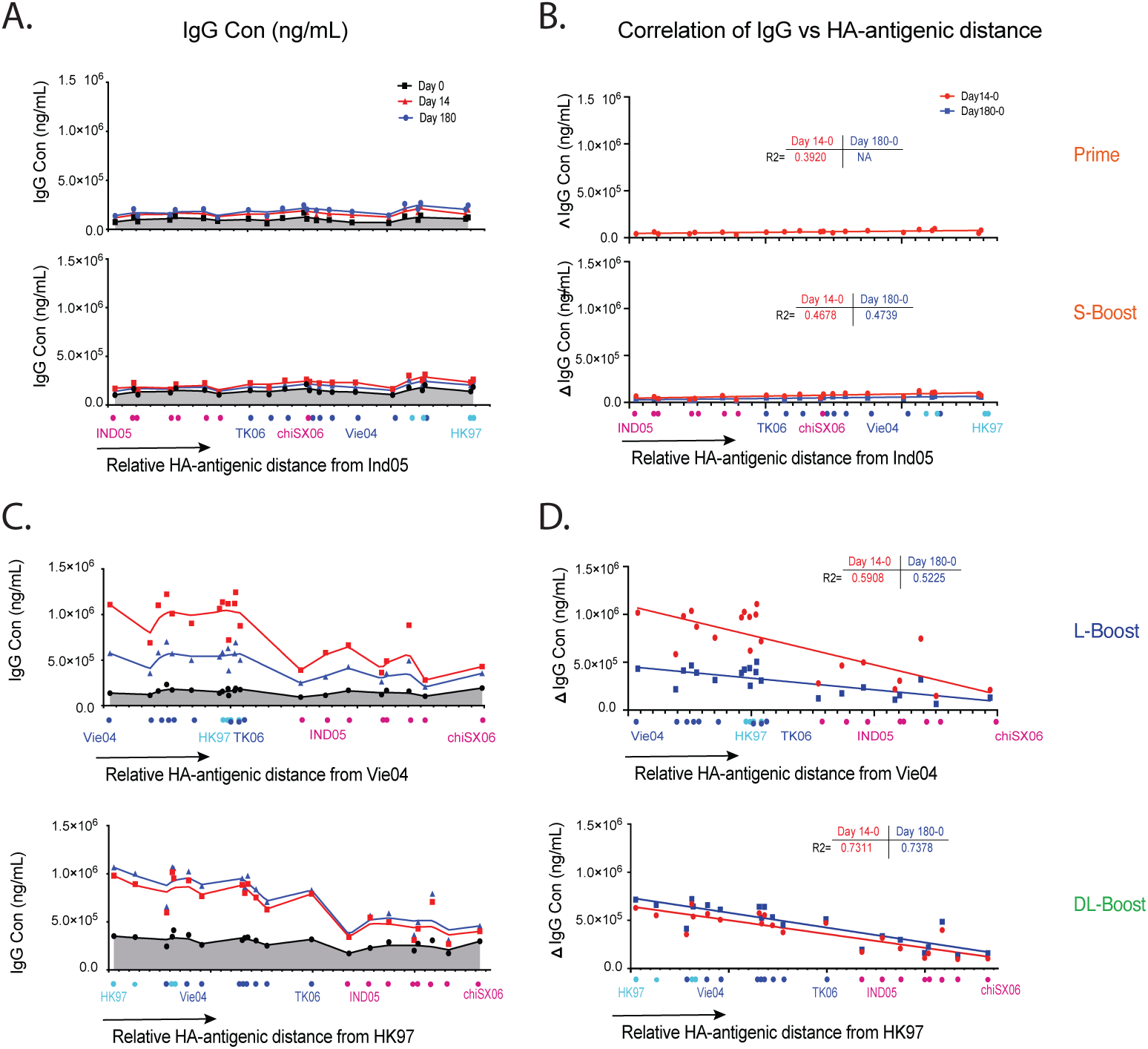
Relative HA antibody landscapes, anti-HA IgG levels and relative antigenic distances from vaccine strains. A. The relative HA antibody landscapes of H5 virus strains as a function of the relative HA antigenic similarity distance from the vaccination strain Ind05 for the Prime group and short interval boost (S-boost) group (see Materials and Methods). B. Correlation of the HA antibody response to the HA-antigenic distance from the vaccine strain HAs of the Prime and S-boost groups. The coordinates of each H5 strain result represent the relative antigenic distance of H5 HA *i* to the vaccine strain HA on each axis. C. Relative HA antibody landscapes for each group using the relative HA antigenic distance from the H5 reference strains A/Vietnam/1203/2004 (Vie04; clade 1), or A/Hong Kong/97 (HK97, clade 0). D. The correlation between the HA antibody response and the HA-antigenic distance of "1rst exposer" H5 strain: Vie04 for the long-interval boost group (L-boost)or HK97 for the double long-interval boost group (DL-boost). The change of IgG concentration (∆*I g G*_*conc*_) is the difference between the anti-HA antibody concentration of past-vaccination from that of prior vaccination. The *R*^2^ values were calculated from linear regression fitting.

We found that the immune response in the Prime and S-boost groups were very weak, and since subjects in these groups were only exposed to the Ind05 MIV strain, we made antigenic landscapes (27) using Ind05 as the reference influenza virus strain. The relative antigenic landscapes for these two groups at days 0, 14 and 180 are shown in FIG4 A and B. Similarly, the serological responses of the L-boost and D-boost groups after boosting were plotted against the antigenic distance relative to Vie04 and HK97, shown in FIG4 C and D. Note that the antigenic distance between the cognate vaccine strain and itself is zero (e.g. Vie04 - Vie04 = 0). The Ind05 MIV showed very low antigenicity in both naive subject groups. Changes in IgG concentration (∆*I g G* = [*I g G*_*t*_] − [*I g G*_*day0*_]) were not correlated with antigenic distance (P = 0.014 and 0.020). However, Ind05 MIV boosting showed higher antibody responses to HAs from strains with a smaller antigenic distance in both L-boost (*R*^2^ = 0.57) and DL-boost groups (*R*^2^ = 0.73). These results support our hypothesis that that the imprinting of primed individuals is highly correlated with the related antigenic distance to the priming strains for long-interval H5 vaccination. FIG 4.

### Long-interval boosting with H5 MIV induced broadly heterosubtypic antibody responses against Group 1 influenza strains

To assess the breadth of heterosubtypic immunity generated by the H5 MIV prime and boost strategy, including IgG reactive against other influenza strain HAs, we estimated antibody cross-reactivity to select group 1 (H1, H2, H5, H6, and H9) and group 2 (H3, H4, H7) HAs (Table S1)) using the mPlex-Flu assay (FIG 5). In all subjects, we detected high pre-existing anti-H1 HA subtype IgG levels against older (A/South Carolina/1/18 (SC18), A/Puerto Rico/8/1934 (PR8)) and newer (A/New Caledonia/20/1999 (NewCall99), A/California/07/2009 (Cali09)) strains. However, these anti-HA levels were not significantly affected by H5 MIV vaccination (FIG S7 A). In addition, we found dramatic increases in anti-HA IgG levels targeting other group 1 influenza viruses (e.g. H2, H6) that had lower baseline levels compared to those against influenza group 2 (H1, H3) subtype HAs.

**FIG 5.**
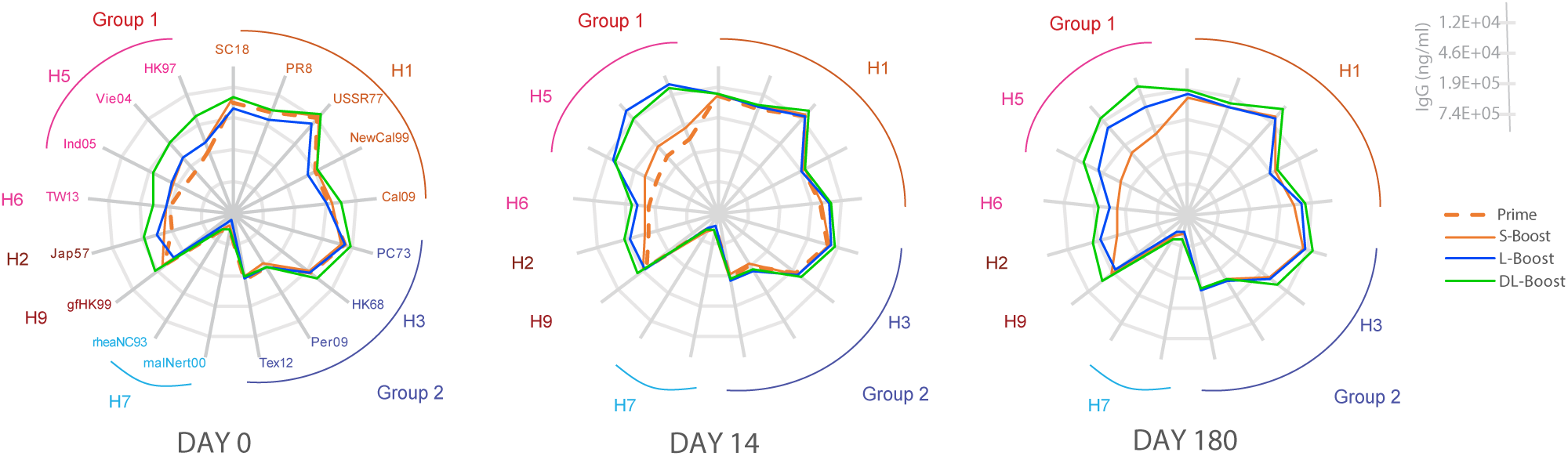
The human heterosubtypic IgG antibody response elicited by H5 MIV. The IgG antibody response induced by H5 influenza vaccine against previously circulating or vaccine influenza strains (H1, H2, H3, H6, H7, H9), were measured by mPlex-Flu assay pre-(day 0) and post-vaccination (days 14, 180).

Further analysis demonstrated that post-H5 vaccination IgG reactivity across influenza strains was inversely correlated to both phylogenetic and antigenic distance between the strains, especially the stalk regions. Based on phylogenetic distance, the gene sequence of H6 is closer to H5 than H9 (33). Similarly, the gene sequence of H2 is closer to H5 than H6 and H1 (FIG S1 A). In addition, we found that IgG responses induced by H5 MIV against HA of A/Japan/305/1957 (Jap57, H2) were significantly higher than that against A/Taiwan/2/2013 (TW13, H6) and A/guinea fowl/Hong Kong/WF10/1999 (gfHK99, H9) (FIG5, FIG S7 A), the latter two strains have stalk regions phylogenetically and antigenically distant from the H5 clade stalk. We also found that, in both primed groups, H5 MIV elicited cross-reactive anti-H2 IgG responses in naive subjects, with a higher peak and a sustained duration than in the unprimed groups. Those responses were stronger than those against H6 and H9 HAs. No significant changes were detected in IgG levels against H3 and other group 2 influenza strains (FIG S7 B). Together, these findings also support the hypothesis that cross-strain, anti-HA antibody responses are highly correlated with phylogenetic similarity, and inversely correlated with antigenic distance, to the vaccine strain.

### Long-interval boosting elicited IgG antibodies against the HA head domain

The HA stalk domain is highly conserved within influenza virus phylogenetic groups, and stalk-reactive antibodies have been hypothesized to be the major contributors mediating cross-reactivity of anti-HA IgG antibodies across group 1(34) strains. However, broadly cross-reactive neutralizing antibodies against the HA head domain have recently been identified, and could also contribute to this phenomenon (reviewed in (35)). Thus, we next measured the change in the relative proportions of head versus stalk reactive IgG within H5 boosting group.

H5 head (HA1) specific IgG levels were measured using beads coupled with the Ind05 head domain only. Anti-stalk IgG was measured using chimeric cH9/1 and cH4/7 proteins to estimate, respectively, group 1 and group 2 stalk-reactive antibodies (36, 37, 38). The results demonstrate that short-interval boosting can induce an ∼2 fold increase in anti-H5 head IgG levels in naive subjects (FIG 6). In addition, significant increases in head-specific IgG were also detected in the L-boost group: 27 fold (14d), 20 fold (28d), and 10 fold (180d). Examining the DL-boost group, ∼7-8 fold increases were observed at 14, 28, 180 days after vaccination. High levels of group 1 stalk-reactive IgG were found in both boosting groups. However, these increases accounted for less than a 2-fold overall change in IgG levels, primarily because these stalk-reactive IgG antibodies were present at relatively high levels prior to vaccination. We did not observe any signi1cant post-vaccination increases in group 2 stalk-reactive antibody levels regardless of test groups. Overall, our results suggest that broadly cross-reactive IgG against H5 influenza HAs or the phylogenetic group 1 are most likely mediated by conserved epitopes on the head domain of HA as opposed to the stalk domain.

**FIG 6.**
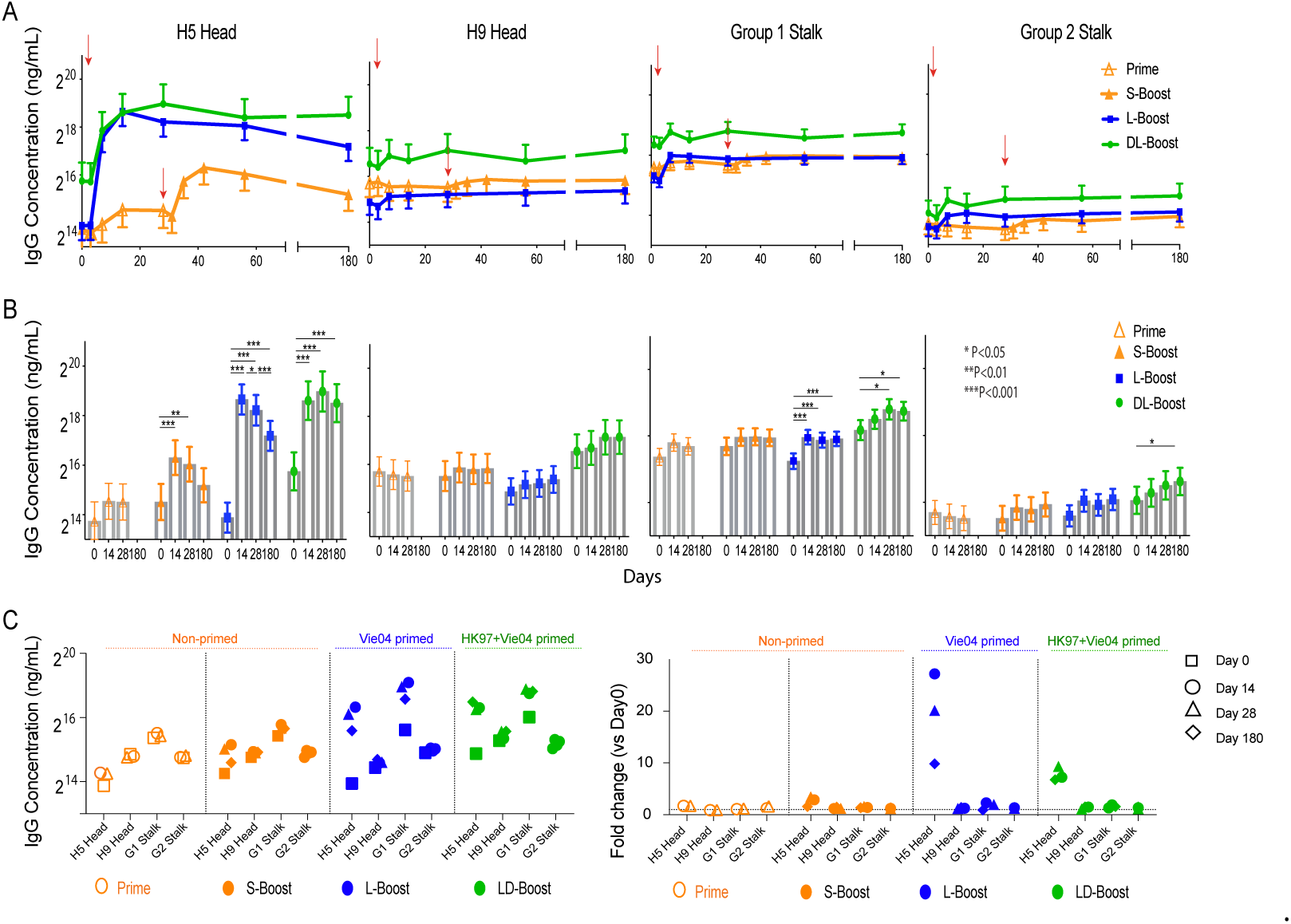
The head and stalk-reactive IgG response induced by the human MIV H5 vaccine. A. The kinetic profile of the IgG response against the HA head or stalk domain estimated by mPlex-Flu assay. B. Comparison of concentrations of each H5 HA speci1c anti-body pre- (day 0) and post-vaccination (14, 28 and 180 days). Linear contrasts within the linear mixed effects models framework were used for statistic testing (^*^ P<0.05, ^**^P<0.01, ^***^P<0.001). C. Comparison of anti-HA IgG concentrations between HAs, including antibodies against chimeric cH9/1 HA (termed group 1 stalk-reactive antibodies; G1 Stalk), and cH4/7 HA (termed group 2 stalk-reactive antibodies; G2 Stalk).

## DISCUSSION

Two major impediments to universal flu vaccine development are the constant antigenic changes of influenza strains, and that the human antibody response is shaped by prior influenza exposure history (39). In addition, vaccination strategies for emergent influenza strains need to take into account both the vaccination schedule, and the ability of HA imprinting to can hinder immune responses to new antigens. Antibody mediated immune responses to new influenza HA antigens are generally weak after the priming vaccination, and require further boosting to elicit adequate titers for infection prevention. This phenomenon can be leveraged if the subject has been primed by exposure to influenza HA antigens, by prior infection or vaccination, that are a short antigenic distance from emergent strain HAs (heterosubtypic immunity).

The antigenic distance between two influenza strain HAs can be calculated empirically or experimentally. Empirically, antigenic distance is the difference between amino acid sequences of HA proteins (e.g. edit distance, Damerau-Levenshtein distance). Experimentally it can be derived by calculating the n-dimensional distance between immune reactivity of sera from a subject vaccinated with a single virus against a panel of other HAs from disparate influenza strains (39). As we have previously shown (37), the smaller the antigenic distance between the prime and boost HAs, the stronger the post-boost vaccination increase in vaccine specific anti-HA IgG levels. This work extends that observation to show that boosting also increases anti-HA IgG to heterosubtypic strains within close antigenic distance of the priming strain.

In this study, we also analyzed changes in multi-dimensional anti-H5 HA IgG responses after vaccination and boosting using a modification of the antibody landscape method (30), a variant of antigenic cartography (32). We initially analyzed anti-HA IgG antibody levels against a comprehensive panel of H5 clade/subclade HAs as a function of the relative antigenic distance to the reference vaccine HA. We call this multi-dimensional measure the *relative antibody landscape* (Fig 4 A and C). This novel method, combined with multiplex serum IgG measurements, allows an analysis of the breadth of the antibody response as a function of the antigenic distance from the vaccine strain. Our results using the relative antibody landscape method show that the anti-H5 HA IgG responses elicited by boosting in both primed groups are highly correlated with the antigenic distance between the priming and boosting H5 vaccine strains. These findings provide further evidence of for the influence HA antigenic imprinting in H5 influenza vaccination. Most significantly, we demonstrate that relative antibody landscape methods can be used to analyze the effects of previous HA antigen exposure on vaccine responses, allowing for quantitative analysis of antigenic imprinting.

Our work also demonstrates that long-interval boosting augments H5 vaccine-induced immunity. Studies using variants of the influenza H5 MIVs have shown that long-interval prime-boost strategies, on the order of 4-8 years between vaccinations, result in robust and durable antibody responses (9) to what are relatively poorly immunogenic vaccine components (11, 12, 24). Intermediate intervals of 6-12 months between priming and boosting with H5 variants significantly increases antibody responses (40, 41), compared to 8 weeks or less. One potential mechanism for these results is a time-dependent increase in long-lived memory B cells, which may take 2-4 months after vaccine priming (42). These memory B cells can then respond rapidly to long interval boosting (43). Significant additional work is necessary to define the optimum prime-boost interval for robust responses.

Our results also support the hypothesis that long-interval boosting increases antibody responses targeting the HA head domain, rather than the stalk. Recently, several broadly neutralizing antibodies (bnAbs) have been identified from both infected or vaccinated human subjects that target the hypervariable HA head domain, including C05 (44), 5J8 (45), CH65 (46) and CH67. These bnAbs exhibit considerable neutralizing breadth within the H1 (44, 45, 46) and H3 (47) influenza subtypes. Such bnAbs are thought to bind highly conserved regions on the sialic acid receptor binding site (RBS) in the HA head domain, explaining their ability to broadly neutralize viral binding from different subtypes (46, 48). As the head domain is known to be immunodominant in the induction of strong antibody responses, broadly head-reactive antibodies could be the major mediator of cross-reactive immunity across influenza subtypes or heterosubtypes. Our results are also consistent with recent work that found rapid activation and expansion of pre-existing memory B cell responses to the conserved epitopes on the HA stalk and head domains after long interval prime-boost vaccination with H7N9 (42).

Finally, our results contribute further to a framework for thinking about universal influenza vaccine development strategies. The aspirational goal of a universal influenza vaccine is to create long-lasting protective immunity to a wide spectrum of influenza viruses. In such cases, future exposure, via infection or vaccination may occur years after the initial priming and imprinting event. Our work demonstrates that the long interval prime-boost strategy for H5 vaccination induces long-lasting cross-reactive antibodies against conserved regions on the HA1 head domain. This may help in universal influenza vaccine development not as a single vaccine, but as a long-interval boost strategy to generate cross-reactive antibodies to recognize the conserved sites on HA1 head domain.

In conclusion, we used a multiplex antibody assay and a novel antibody landscape method to analyze antibody mediated immunity to various influenza HAs after H5 vaccine priming and boosting. These methods quantitatively account for the antigenic distances between the vaccine and other strain HAs. This new approach demonstrated that anti-H5 IgG antibody responses elicited by boosting are highly correlated to the antigenic distance between the the priming and boosting H5 vaccine strains, providing evidence for OAS and HA imprinting within the context of H5 vaccination.

## MATERIALS AND METHODS

### Human Subjects Ethics Statement

This sub-analysis study was approved by the Research Subjects Review Board at the University of Rochester Medical Center (RSRB approval number RSRB00012232). Samples were analyzed under secondary use consent obtained previously as part of prior clinical trial (24). All research data were coded by sample IDs in compliance with the Department of Health and Human Services’ Regulations for the Protection of Human Subjects (45 CFR 46.101(b)(4)).

### Samples and data

Serum samples for the multiplex assay were obtained from a prior clinical trial, DMID 08-0059 (Figure 1)(24). Subjects without pre-vaccination serum samples (Day 0 baseline) were excluded. All subjects in the three cohorts were inoculated with inactivated A/Indonesia/5/05 (A/Ind05) vaccine. H5 naive subjects (*n* = 12), who were healthy adults, not at risk for H5 exposure and with no H5 vaccination history, received 2 identical A/Ind05 vaccinations separated by 28 days. Primed subjects (*n* = 30) previously received the inactivated subvirion influenza A/Vietnam/1203/04 (A/Vie04) vaccine in 2005–2006 (9). The double primed group (*n* = 13) had received both the recombinant influenza A/Hong Kong/156/97 vaccine (A/HK97) in 1997-1998 (11) and the influenza A/Vie04 vaccine in 2005-2006. Serum samples were collected before vaccination (Day 0) and on days 7, 14, 28, 56, and 180 after vaccination. Serum samples were collected from the naive group subjects on days 7, 14, and 28 days after the second immunization. All data from the mPlex-Flu, HAI, and MN assays were adjusted for dose difference using linear mixed effects models, as previously described (28, 29).

### mPLEX-Flu Analysis

We estimated the concentrations of anti-HA IgG antibodies against a 45 HA antigen panel of influenza viruses using the mPLEX-Flu assay, as described previously(25, 36). All influenza HA sequence identi1ers uesd are listed in the TABLE S1 and the HA genetic distance (phylogenic tree) is shown in FIG S1 A. The panel recombinant HA proteins were expressed by baculovirus system and puri1ed Ni^+^ affinity column selection as previously described (36) and verified (FIG S1 B.

The calculation of individual IgG concentrations for each influenza strain anti-HA IgG was performed using standard curves generated from 1ve-parameter logistic regression models (28, 29). All IgG concentration results from the mPlex-Flu assay was adjusted using linear mixed effects models accounting for the group, day, and group-day interactions for each H5 vaccine strain. Covariates adjusted in the linear mixed effects models included age at enrollment, gender, ethnicity (Caucasian vs. non-Caucasian), dose (two dose levels: 15 and 90 *µg*), and analytic batch (five batches) factors (28, 29).

### Antigenic cartography of H5 influenza viruses generated by mPlex-Flu assay data

In order to estimate the antigenic distance of HA antigens of H5 influenza virus strains, we adopted the 17 H5 HA genes that covered all 10 clades/subclades strains of H5 from Dr. Paul Zhou from Institute Pasteur of Shanghai, Chinese Academy of Sciences, Shanghai, China (3). The 17 individual antisera against each H5 influenza virus strain were generated with mouse DNA vaccination as previously described (3), and shown in FIG S2 A. Using the mPlex-Flu assay, we evaluated the 17 anti-sera against a panel of 36 HA antigens to create a multi-dimensional matrix, after normalizing the dilution factors and subtracting the background levels, using generalized linear models with identity link functions (FIG S2 B). Classical multidimensional scaling was used to project multi-dimensional distances into two-dimensional antigenic cartography plots plots(31, 25). The coordinates for two-dimension antigenic cartography were further used to calculate the Euclidean distance between H5 influenza viruses to obtain the antigenic distance matrix(FIG S3).

### Relative antigenic landscapes of antibody response

Based on the antigenic distances generated above, and using the three vaccine strains as reference: A/Hong Kong/156/97 vaccine (HK97, clade 0) A/Vietnam/1203/04 (Vie04, clade 1) A/Indonesia/5/05 (Ind05, clade 2) a vaccine-strain relative antigenic distance matrix was selected. Next, relative antigenic antibody landscape-like figures were created by using the relative antigenic distance as the X-axis and the Y-axis is IgG antibody response. Data points were linked by LOWESS 1t spline curves (Prism 8 software). A set of antibody response landscape-like plots were generated for each vaccination strain.

### H5 head and stalk speci1c antibody response

We used the mPlex-Flu assay to simultaneously assess the antibodies to the head and stalk domains of HA. We coupled Luminex beads with the head region of HA, which are puri1ed recommbinant proteins of HA1 domain of H5/Ind05 and H9/A/guinea fowl/Hong Kong/WF10/1999 (gfHK99, H9). To detect the group 1 stalk-reactive antibodies, we used the chimeric cH5/H1 (head/stalk) and cH9/H1 proteins. For group 2 stalk-reactive antibodies, we used the cH5/H3 and cH7/H4 proteins kindly provided by Dr. Florian Krammer(49, 34, 36, 37).

### Reanalyses of HAI and MN data

Primary HAI and MN data were generated previously during the vaccine trial as described (24). Serum antibody responses to the homologous A/Indonesia/05/2005 PR8-IBCDC-RG2 virus were measured at the Southern Research Institute (11). We reanalyzed these data using linear mixed effects models, with correlations from repeated measurements within the same subject considered. The same predictors and covariates were used in the linear mixed effects models for the HAI and MN data analysis as for the mPLEX-Flu data analysis (28).

### Availability of data and materials

All data generated in this study are included in this published article and in the Supplementary Material.

## ACKNOWLEDGMENTS

We would like to thank Dr. Paul Zhou from Institute Pasteur of Shanghai, Chinese Academy of Sciences, Shanghai, China for providing the H5 HA constructs used to generate the mouse anti-sera for antigenic cartography, and Dr. Florian Kramer, Ichan School of Medicine at Mount Sinai, New York, United States for several influenza single strain and chimeric HA constructs.

This work was supported by the National Institutes of Health Institute of Allergy, Immunology and Infectious Diseases grants R21 AI138500 (MZ, JW, AW, SP), R01 AI129518 (MZ, SPH, JW, AW, SP) and the University of Rochester Clinical and Translational Science Award UL1 TR002001 from the National Center for Advancing Translational Sciences of the National Institutes of Health (JW, DL, MZ). The content is solely the responsibility of the authors and does not necessarily represent the oZcial views of the National Institutes of Health.

J.W. and M.Z. conceived of the project, designed and oversaw the experiments the experiments and analysis, and wrote the paper. J.T. provided the data and samples from the prior DMID 08-0059 study, and contributed to the study design. D.L. performed the statistical analyses and modeling. J.W. S.P. and A.W performed the experiments. S.P.H. contributed to the experimental design and wrote the paper. M.S. contributed substantially to the design and analysis of the imprinting experiments. All authors read and approved the manuscript.

We declare no competing interests.

## Supplementary Material

Generation of an antigenic cartography representing 21 clades or subclades of H5 influenza viruses using mouse antisera reactivity measured using our mPlex-Flu assay

### 1. ANIMALS

Female BALB/c mice were purchased from Taconic Biosciences. For all experiments, female 8- to 12-week-old mice were used and randomly assigned to experimental groups. All research involving live, vertebrate animals was conducted in accordance with the Public Health Service Policy on Human Care and Use of Laboratory Animals. mice were maintained at the University of Rochester Medical Center Vivarium, a AAALAC certified Vivarium (Animal Welfare Assurance Number is A-3292-01), under their established guidelines, including isolation, feeding, recovery procedures, and euthanasia in accordance with Federal regulations. All experimental procedures for animals were approved by the Institutional Animal Care and Use Committee (IUCAC; protocol number UCAR-2011-055E), and all personnel working with the animals were trained and certified by the IUCAC and Vivarium staff.

### 2. GENERATION OF MOUSE ANTISERA DIRECTED AGAINST HEMAGGLUTININS (HAS) REPRESENTING H5 CLADES AND SUBCLADES

A panel of 17 total DNA plasmids encoding all 10 H5 clades/subclades HA genes was provided by Dr. Paul Zhou from Institute Pasteur of Shanghai, Chinese Academy of Sciences, Shanghai, China (Zhou et al., 2012). The DNA vaccination plamids were constructed from the mammalian expression vector, pCMV/R, containing whole codon-optimized HA gene inserts representing H5 influenza viruses (Zhou et al., 2012). All above plamid DNAs were amplified and then purified with Plasmid Maxi Kit (QIAGEN) following the manufacture’s recommendations. Purified DNA plamids were used for intramuscular immunizations (i.m.) of 8- to 12-week-old BALB/c mice as previously described (Zhou et al., 2012). Briefly, the mice (n=4) were inoculated with 100g of one of the 17 H5 subclade HA plamid DNAs respectively, on days 0, 28 and 56 (see Figure S2 A). Fourteen days after the last immunization, serum samples were collected from each mouse and combined within each group. The resultant H5 reactive antisera panel was aliquoted and stored at −20^◦^C for future analysis.

### 3. RECOMBINANT HA PROTEINS (RHAS) OF H5 CLADES AND SUBCLADES

All rHAs of type A and B influenza viruses used in this study (TABLE S1) were expressed using pFastBac baculovirus system with a C-terminal trimerization domain and a hexahistidine purification tag (Wang et al., 2018). The entire panel of 17 H5 clades and subclades HA genes were subcloned into this pFastBac vector using BamHI and NotI restriction endonucleases (NEB, Ipswich, MA) as previously published (Wang et al., 2018). We also synthesized four HA genes (see TABLE S1) of H5 avian influenza viruses belonging to the new subclade 2.3.4.4 that circulated during the huge H5 outbreak in turkey and chicken farms in USA in 2015, including A/duck/Sichuan/NCXN10/2014 (ducSC14,gene bank accession No:KM251469), A/turkey/Washington/61-22/2014 (turWash14,accession No:KP739397), A/duck/Guangdong/wy11/2008 (ducGD08, accession No:CY091627) and A/turkey/California/K1500169-1.2/2015 (turCal15, accession No: KR150901). We also inserted these HA genes into the pFastBac expression vector to express the rHAs of H5 influenza virus HAs.

Expression and purification of rHA was performed as previously described (Wang et al., 2018). Purified rHAs were concentrated and desalted with 30 kDa Amicon Ultracell centrifugation units (Millipore, Billerica, MA) and re-suspended in phosphate buffered saline (PBS, pH7.4). The purity, integrity and identity of proteins was assessed by NuPage 4-12% Bis-Tris gels (Invitrogen, Grand Island, NY), the results of which are shown in FIG S1 B. Protein concentration was quantified using the Quickstart Bradford Dye Reagent (Bio-Rad, Hercules, CA) with a bovine serum albumin standard curve.

### 4. SIZE EXCLUSION CHROMATOGRAPHY (SEC) ANALYSIS OF RHAS OF H5 INFLUENZA VIRUSES

The four representative rHAs of H5 influenza viruses were evaluated, including A/Chicken/Guangxi/12/2004 (chiGX04, CL2.4), A/Silky Chicken/Hong Kong/SF189/01 (s.chiHK01, CL3), A/Goose/Guiyang/337/2006 (gooGY06, CL4) and A/Chicken/Shanxi/2/2006 (chiSX06, CL7.2) (Colored in FIG S3. A). Partially cleaved HA0 (into HA1 and HA2) and uncleaved HA0 were analyzed by SEC using the AKTA chromatography system (GE Healthcare Bio-Sciences, Pittsburgh, PA) through a HPLC Biosep-SEC-s4000 column (300×7.8mm, 00H-2147-K0, Phenomenex Inc, Torrance, CA) in pH 6.8 buffer of 50mM Na2HPO4, 50mM NaHPO4 and 150mM NaCl at 1.0 ml/min flow rate. All four purified HA preps took the same length of time to flow out from SEC column in HPLC analysis (see FIG S1 C), suggesting that there are not significant differences between the sizes of the protein preps, regardless of whether the HA is uncleaved, partially or fully cleaved. This result provides evidence that the cleavage of HA0 does not affect HA1 and HA2 binding to form the trimmer structure in the natural condition. The protein standards for SEC were purchased from Bio-Rad (151-1901, Bio-Rad Inc, Hercules, CA), and the peaks of protein standards in HPLC analysis are shown in FIG S1 C.

### 5. DEVELOPMENT OF H5 MPLEX-FLU ASSAY

The mPlex-Flu assay contained an HA antigen panel with 35 rHAs from various influenza strains, HA domains and chimeric HAs (Supplementary Table S1). The phylogenetic amino acid sequence tree of those 35 HA proteins is shown in FigureS1.

We used the mPlex-Flu assay to estimate the strain specific binding of the 17 H5 DNA vaccination anti-sera with our 35 HA antigen panel as previously described (Wang et al., 2018). After normalizing the dilution factor using the generalized linear model with identity link function, strain specific binding was obtained from the estimated coefficients, using the background binding as the reference group (i.e. the strain specific binding is equal to the estimated strain binding minus the background binding).

### 6. GENERATION AN ANTIGENIC CARTOGRAM UTILIZING A NOVEL MULTIPLE DIMENSIONAL SCALING METHOD (MDS)

Multidimensional scaling preserves the dissimilarities among strains by generating a two-dimension plot with the distance between each strain approximating their dissimilarities. Using classical (metric) MDS, multiple dimensional matrix data were generated using mPlex-Flu assay data, as shown in Figure S2, B. The resultant two-dimensional MDS plot allowed us to estimate the H5 HA antigenic distances and generate the HA antigenic cartography as previously published (Zhou et al., 2012; Smith et al., 2004). Due to the continuous nature of data from the mPlex-Flu assay (as compared to the relative discrete data from the HAI assay) and the consistent range of estimated strain specific binding, we constructed the antigenic map by minimizing different error functions *E* = Σ_*ij*_*e*(*MFI*_*ij*_,*d*_*ij*_, where *MFI*_*ij*_ denotes the target distance between antigen *i* and antiserum *j* and *dij* denotes the Euclidean distance between antigen *i* and antiserum *j* in the two-dimensional map. *e*(*MFI*_*ij*_, *d*_*ij*_) = (*MFI*_*ij*_ − *d*_*ij*_)^2^ is the error function we minimized to construct the antigenic cartogram.

We also conducted sensitivity analysis to compare the generated antigenic cartography with previous published methods (Smith et al., 2004). We replaced 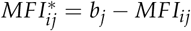 with *MFI*_*ij*_ as input for the MDS method, where *bj* denotes the maximum measurement for antiserum *j*. We obtained the same antigenic cartography as expected due to the consistent range of the estimated strain specific binding, shown in Figure S2, C.

An HA antigenic cartography consisting of 21 H5 influenza virus strains was generated utilizing antisera from H5 DNA vaccinated mice, to calculate the antigenic distance from the H5 A/Hong Kong/1997 clade 0 (HK97(0)), see Supplementary Figure S3.

---

**Fig. S1.**
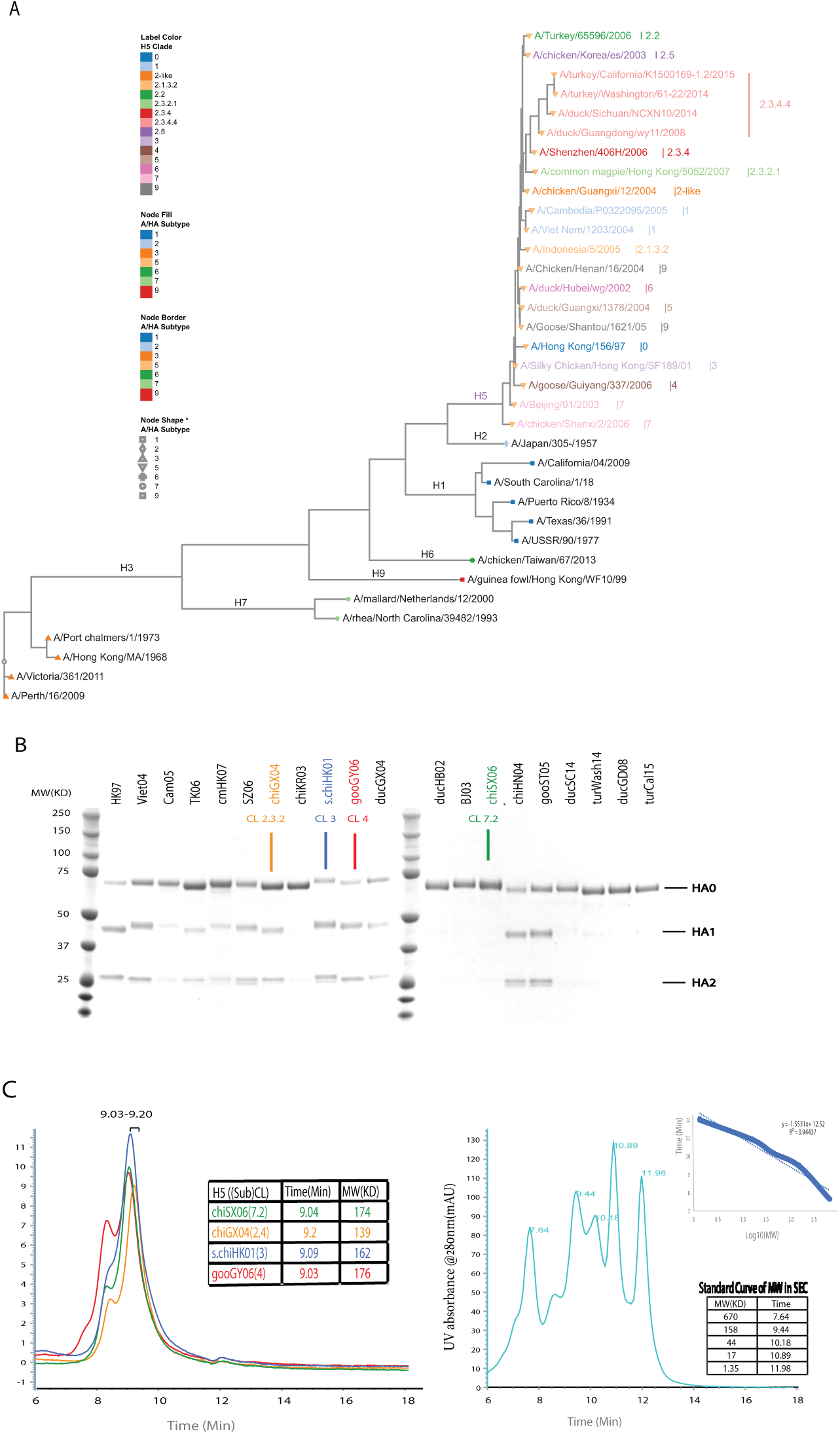
HA protein characters of 35 influenza virus A strains in mPlex-Flu assay. A. The phylogenetic tree was generated using HA amino acid sequences of the 35 influenza A virus strains obtained from the phylogenic tree maker on the Influenza Research Database Website (https://www.fludb.org/brc/home.spg?decorator=influenza). B.SDS-PAGE gel image of purified HA proteins of H5 influenza viral strains. C. HPLC analysis results of four representative HA proteins flowing through the Biosep-SEC-s4000 columns with the Bio-rad protein standards.

**Fig. S2.**
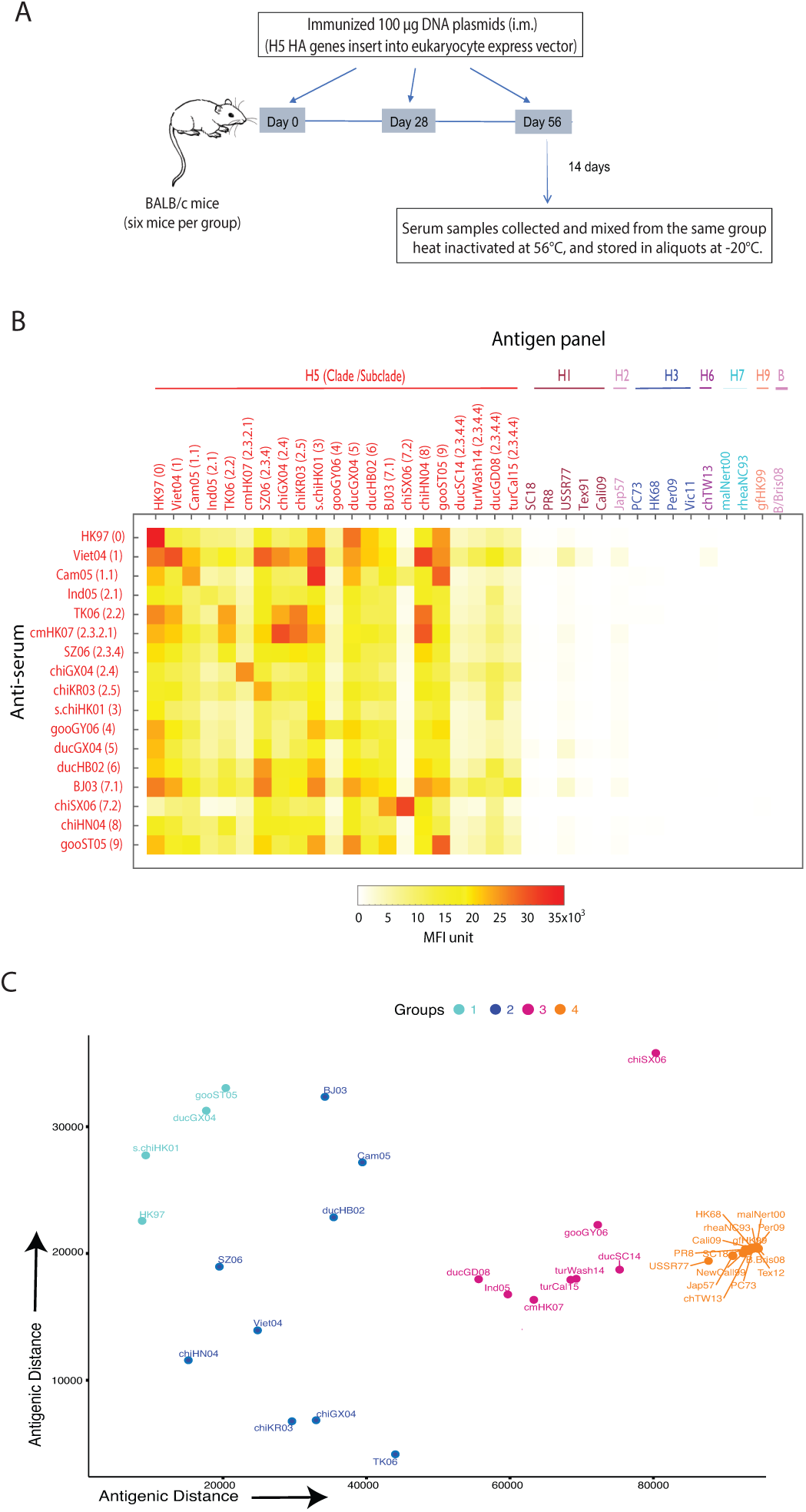
Antigenic cartography is generated with a mouse DNA vaccination model. A. Mouse DNA vaccination strategy. B. Heat map of the multiple dimensional antibody data generated by the mPlex-Flu assay. Each mouse polyclonal antiserum was induced by DNA vaccination with a DNA plasmid encoding HA proteins, and the antibody levels in the sera were estimated by mPlex-Flu assay. C. Antigenic cartography of 36 influenza A strains assessed by mPlex-Flu assay with the Multiple Dimensional Scaling (MDS) method.

**Fig. S3.**
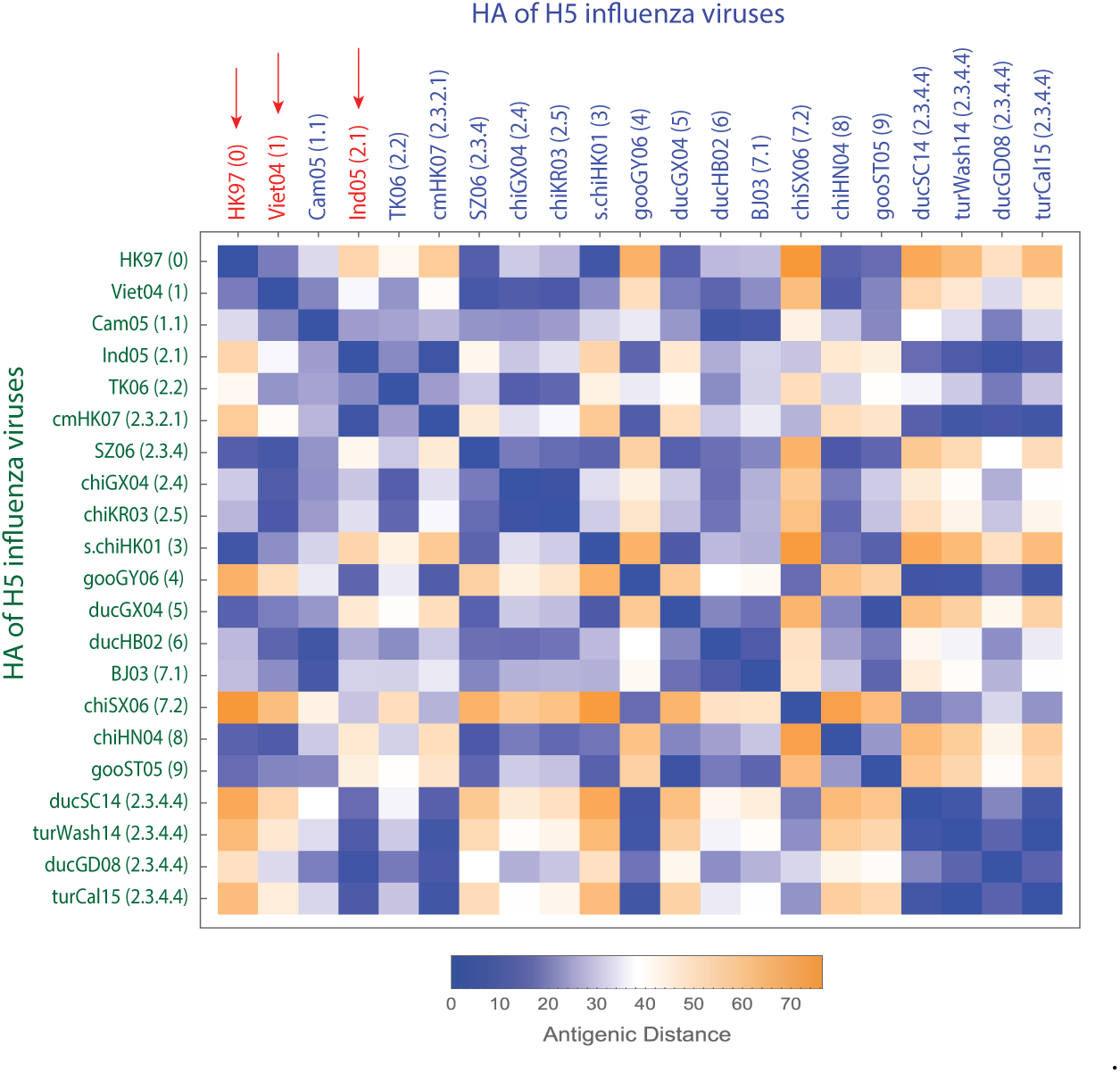
The heat-map matrix of the antigenic distance between the 21 H5 influenza virus strains. The three vaccination strains are highlighted with red arrows.

**Fig. S4.**
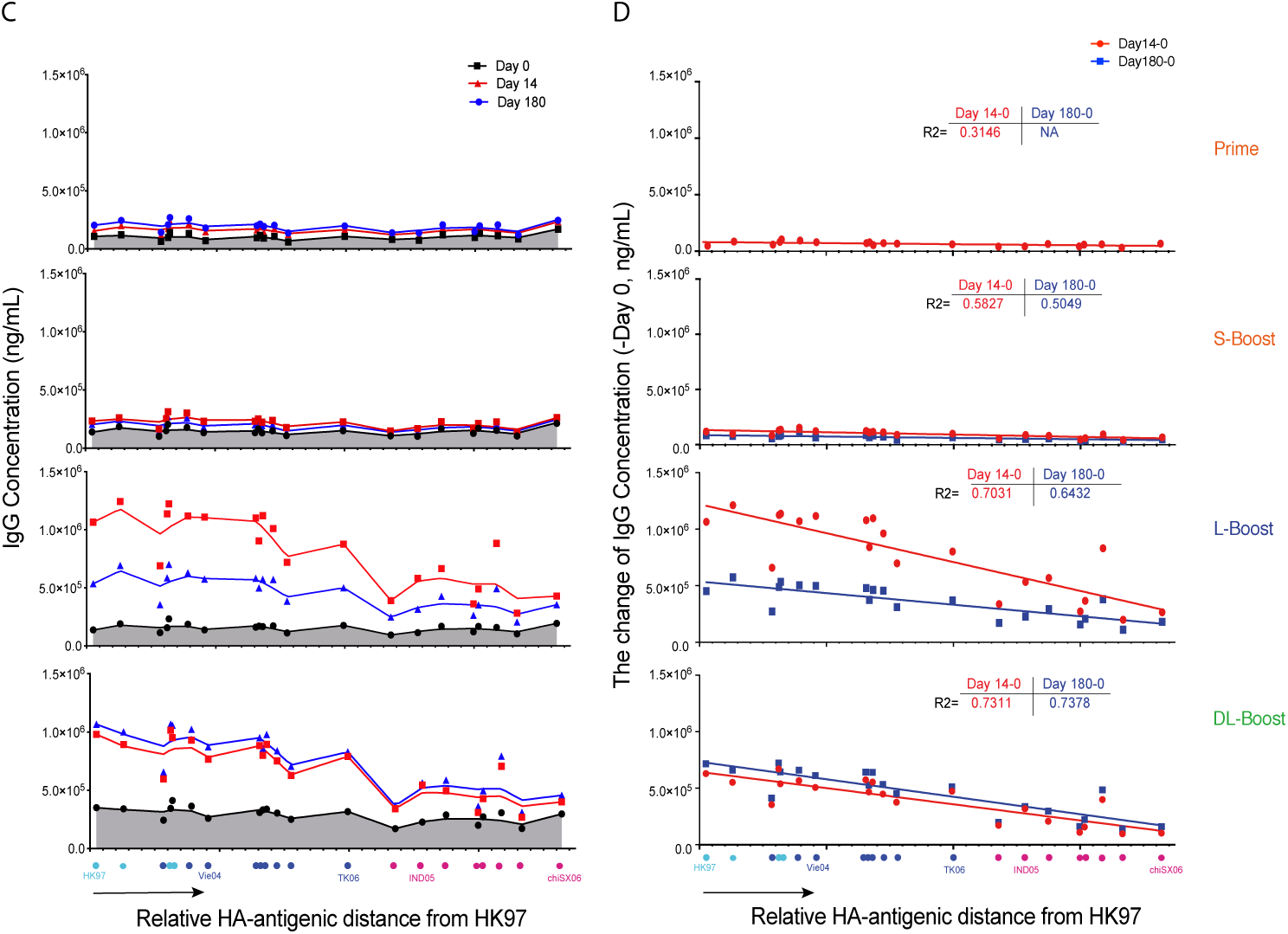
The correlation between the HA antibody response and HA antigenic similarity of A/Hong Kong/156/97 (HK97) to 21 H5 influenza virus strains. A. The HA antibody response landscape-like plots of each group using the relative HA antigenic distance of A/Hong Kong/156/97 (HK97, clade 0) as the reference strains (see material and methods). X-axis is relative antigenic distance; Y-axis is IgG antibody response; the spots were linked by LOWESS fit spline curve (Prism 8 software). B. The correlation of the HA antibody response to the HA-antigenic distance. The ∆ change of antibody concentration of pre- and post-vaccination versus the relative HA antigenic distance of Vie04. The R squared values were calculated with simple linear regression analysis (Prism 8 software).

**Fig. S5.**
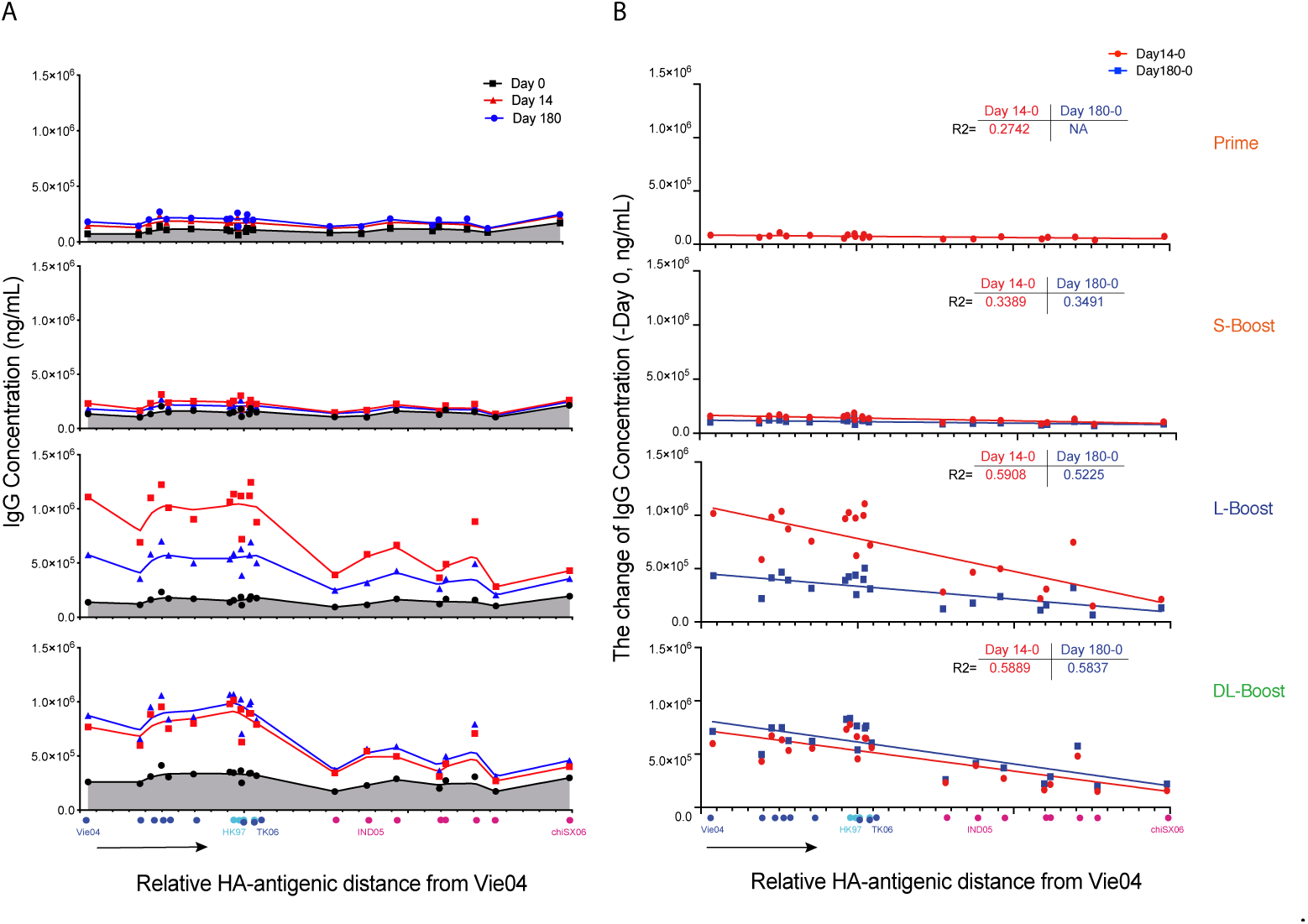
The correlation between the HA antibody response and HA antigenic similarity between A/Vietnam/1203/2004 (Vie04) and 21 H5 influenza virus strains. A. The HA antibody response landscape-like plots of each group using the relative HA antigenic distance of A/Vietnam/1203/2004 (Vie04, clade 1) as the reference strains (see material and methods). X-axis is relative antigenic distance; Y-axis is IgG antibody response; the spots were linked by LOWESS fit spline curve (Prism 8 software). B. The correlation of the HA antibody response to the HA-antigenic distance. The ∆ change of antibody concentration of pre- and post-vaccination versus the relative HA antigenic distance of Vie04. The R squared values were calculated with simple linear regression analysis (Prism 8 software).

**Fig. S6.**
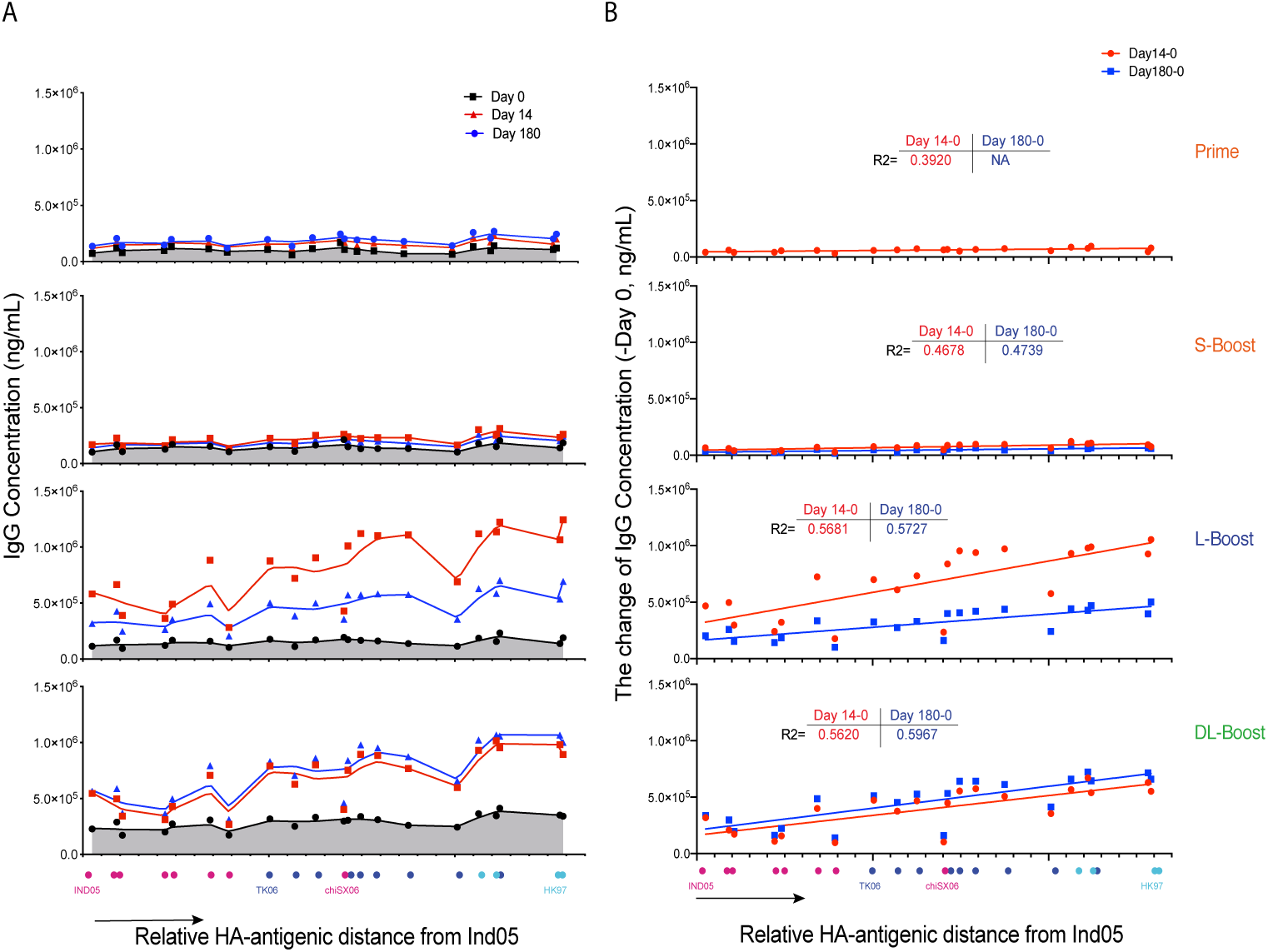
The correlation between the HA antibody response and HA antigenic similarity of A/Indonesia/5/05 (Ind05) to 21 H5 influenza virus strains. A. The HA antibody response landscape-like plots of each group using the relative HA antigenic distance of A/Indonesia/5/05 (Ind05, clade 1) as the reference strains (see material and methods). X-axis is relative antigenic distance; Y-axis is IgG antibody response; the spots were linked by LOWESS fit spline curve (Prism 8 software). B. The correlation of the HA antibody response to the HA-antigenic distance. The ∆ change of antibody concentration of pre- and post-vaccination verse the relative HA antigenic distance of Vie04. The R squared values were calculated with simple linear regression analysis (Prism 8 software).

**Fig. S7.**
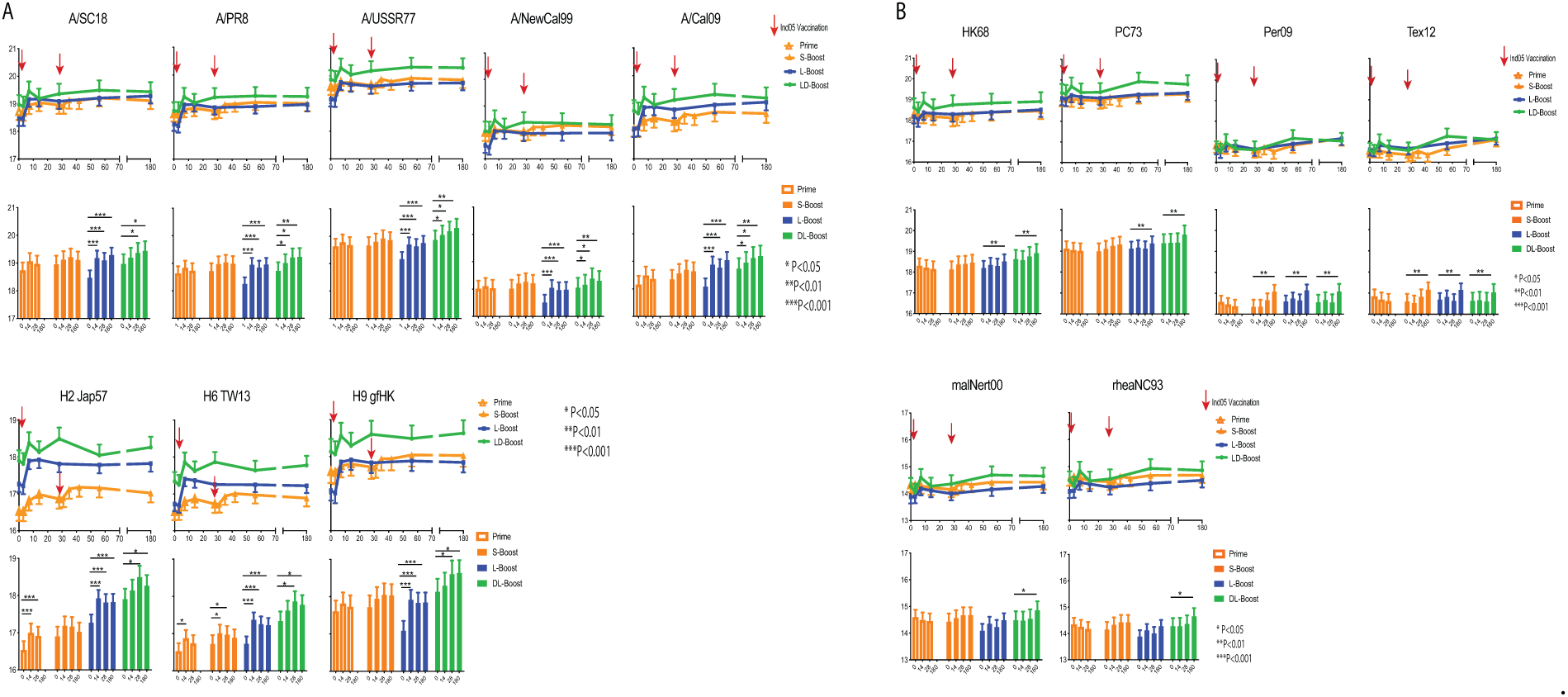
The IgG concentration of group 1 and 2 influenza virus strains was estimated by mPlex-Flu assay in the DMID 08-0059 study. The mPlex-Flu assay estimated the mean and standard deviation of IgG concentration for each group. Then the antibody concentrations were adjusted within the linear mixed-effects models, which included the following: age at enrollment, gender, ethnicity (Caucasian vs. non-Caucasian), dose (two dose levels: 15 and 90 *µg*), and batch (five batches). A. The mPlex-Flu assay estimated the antibody concentrations of group 1 influenza virus strains (including five human H1, one of each H2, H6, and H9). B. The antibody concentrations to group 2 influenza A virus strains (including four H3, and two H7 strains) were estimated by the mPlex-Flu assay.

**Fig. S8.**
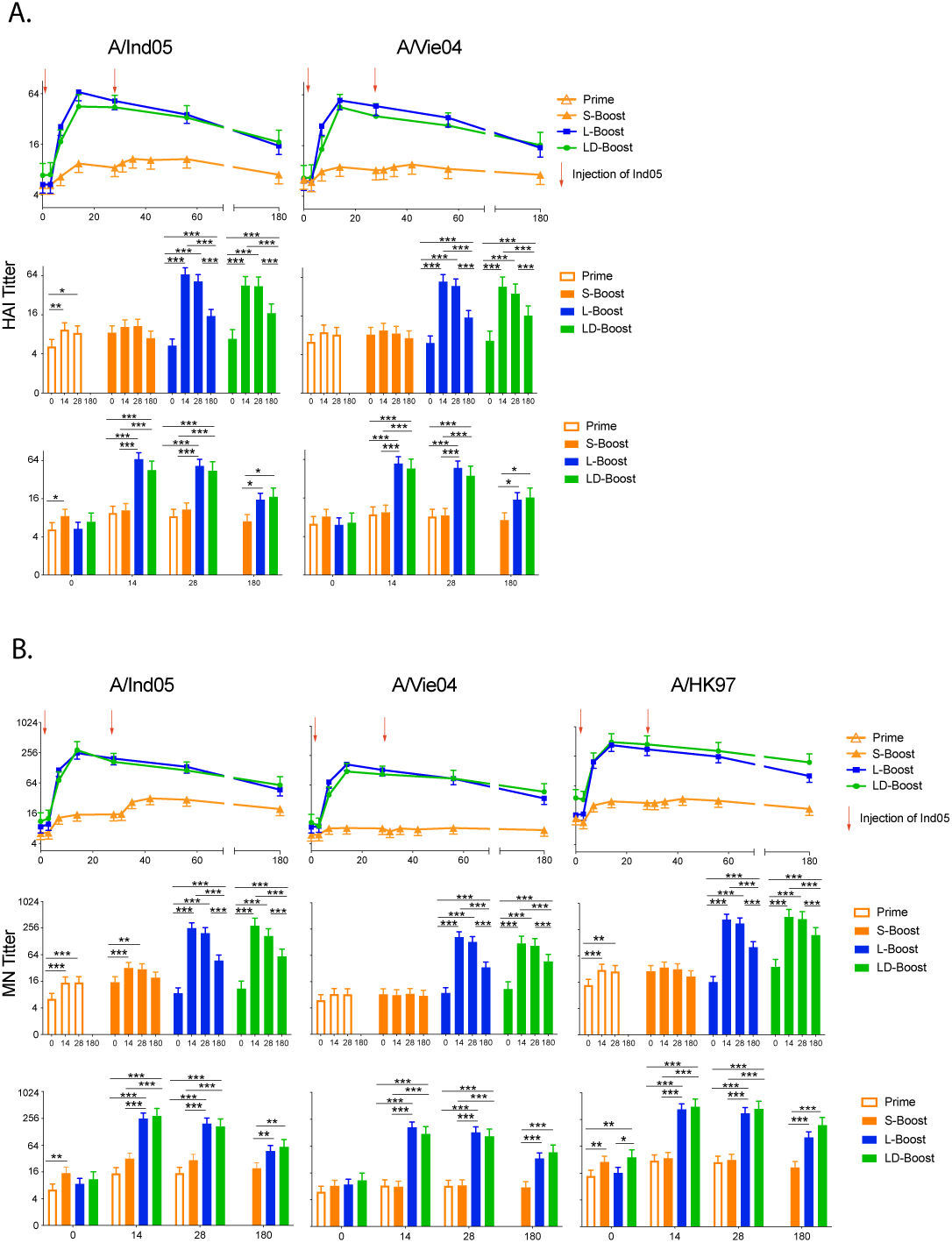
Prior vaccination with a monovalent influenza vaccine (MIV) increased the serum titers of hemagglutination-inhibition (HAI) and micro-neutralization (MN) antibody responses against three antigenically drifted virus vaccine strains, including new vaccine strain A/Indonesia/05/2005 (Ind05; clade 2), previous MIV strains A/Vietnam/1203/2004 (Vie04; clade 1), A/Hong Kong/156/1997 (HK97; clade 0). Naive subjects (Unprimed) received the MIV Ind05 strain and were subsequently boosted at day 28 with the same strain. A previous primed group, received the MIV Vie04 5 years prior, (Primed) then received a single dose of Ind05. The previous double primed MIV Vie04 and HK97 (Multiple). The mean and standard deviation of IgG concentration for each group were estimated by linear mixed effects models with group, day, and group-day interaction used to fit the data for each H5 vaccine strain. Co-variates adjusted in the linear mixed effects models included the following: age at enrollment, gender, ethnicity (Caucasian vs. non-Caucasian), dose (two dose levels: 15 and 90 *µg*), and batch (five batches). * P<0.05, **P<0.01, ***P<0.001 Linear contrasts within the linear mixed effects models framework were used to do the statistical testing.

**Table S1.** The mPlex-Flu assay panel of seasonal influenza viruses, H5 clades and subclades

| Influenza<br>Virus Type | Subtypes | Full Name of Viruses | Abbreviation | H5 Clades<br>/Subclades | Genbank Accession # |
| --- | --- | --- | --- | --- | --- |
| A | H1 | A/South Carolina/1/18 | SC18 |  | AF117241.1 |
|  |  | A/Puerto Rico/8/1934 | PR8 |  | CY148243.1 |
|  |  | A/USSR/90/1977 | USSR77 |  | DQ508897.1 |
|  |  | A/New Caledonia/20/1999 | NewCall99 |  | DQ508889.1 |
|  |  | A/Texas/36/1991 | Tex91 |  | CY125100.1 |
|  |  | A/California/07/2009 | Cali09 |  | FJ966974.1 |
|  | H2 | A/Japan/305/1957 | Jap57 |  | L20407.1 |
|  | H3 | A/Port Chalmers/1/1973 | PC73 |  | CY112249.1 |
|  |  | A/Hong Kong/1/1968 | HK68 |  | CY009348.1 |
|  |  | A/Perth/16/2009 | Per09 |  | GQ293081.1 |
|  |  | A/Victoria/361/2011 | Vic11 |  | KM821347 |
|  |  | A/Texas/50/2012 | Tex12 |  | KC892248.1 |
|  | H5 | A/Hong Kong/156/97 | HK97 | 0 | AF028709 |
|  |  | A/Viet Nam/1203/2004 | Viet04 | 1 | EF541403 |
|  |  | A/Cambodia/P0322095/2005 | Cam05 | 1.1 | HQ200458 |
|  |  | A/Indonesia/5/05 | Ind05 | 2.1.3.2 | EF541394 |
|  |  | A/Turkey/65596/2006 | TK06 | 2.2.1 | EF619998 |
|  |  | A/Common Magpie/Hong Kong/5052/2007 | cmHK07 | 2.3.2.1 | CY036173 |
|  |  | A/Shenzhen/406H/2006 | SZ06 | 2.3.4 | EF137706 |
|  |  | A/Chicken/Guangxi/12/2004 | chiGX04 | 2.4 | DQ366330 |
|  |  | A/Chicken/Korea/es/2003 | chiKR03 | 2.5 | EF541412 |
|  |  | A/Silky Chicken/Hong Kong/SF189/01 | s.chiHK01 | 3 | AF509021 |
|  |  | A/Goose/Guiyang/337/2006 | gooGY06 | 4 | DQ992765 |
|  |  | A/Duck/Guangxi/1378/2004 | ducGX04 | 5 | DQ320884 |
|  |  | A/Duck/Hubei/wg/2002 | ducHB02 | 6 | DQ997094 |
|  |  | A/Beijing/01/2003 | BJ03 | 7.1 | EF587277 |
|  |  | A/Chicken/Shanxi/2/2006 | chiSX06 | 7.2 | DQ914814 |
|  |  | A/Chicken/Henan/16/2004 | chiHN04 | 8 | AY950234 |
|  |  | A/Goose/Shantou/1621/05 | gooST05 | 9 | DQ095628 |
|  |  | A/duck/Sichuan/NCXN10/2014 | ducSC14 | 2.3.4.4 | KM251469 |
|  |  | A/turkey/Washington/61-22/2014 | turWash14 | 2.3.4.4 | KP739397 |
|  |  | A/duck/Guangdong/wy11/2008 | ducGD08 | 2.3.4.4 | CY091627 |
|  |  | A/turkey/California/K1500169-1.2/2015 | turCal15 | 2.3.4.4 | KR150901 |
|  | H6 | A/Taiwan/2/2013 | TW13 |  | KJ162860.1 |
|  | H7 | A/mallard/Netherlands/12/2000 | malNert00 |  | EF470586 |
|  |  | A/rhea/North Carolina/39482/1993 | rheaNC93 |  | KF695239 |
|  | H9 | A/guinea fowl/Hong Kong/WF10/1999 | gfHK99 |  | AY206676.1 |
| HA domains |  | Head of A/Indonesia/5/05 | H5 Head |  |  |
|  |  | Head of A/guinea fowl/Hong Kong/WF10/1999 | H9 head |  |  |
| Chimeric HA |  | cH5/1 (A/Indonesia/5/05, A/Puerto Rico/8/1934) | cH5/1PR |  |  |
|  |  | cH5/1 (A/Indonesia/5/05, A/California/07/2009) | cH5/1Cal |  |  |
|  |  | cH9/1 (A/gf/HK/WF10/1999, A/California/07/2009) | cH9/1 |  |  |
|  |  | cH4/7 (A/duck/Czech/1956, A/Shanghai/1/2013) | cH4/7 |  |  |

